# Diffuse X-ray Scattering from Correlated Motions in a Protein Crystal

**DOI:** 10.1101/805424

**Authors:** Steve P. Meisburger, David A. Case, Nozomi Ando

**Affiliations:** Department of Chemistry, Princeton University, Princeton, New Jersey 08544, USA; Department of Chemistry and Chemical Biology, Cornell University, Ithaca, New York 14850, USA; Department of Chemistry and Chemical Biology, Rutgers University, Piscataway, New Jersey 08854, USA

## Abstract

Protein dynamics are integral to biological function, yet few techniques are sensitive to collective atomic motions. A long-standing goal of X-ray crystallography has been to combine structural information from Bragg diffraction with dynamic information contained in the diffuse scattering background. However, the origin of macromolecular diffuse scattering has been poorly understood, limiting its applicability. We present a detailed diffuse scattering map from triclinic lysozyme that resolves both inter- and intramolecular correlations. These correlations are studied theoretically using both all-atom molecular dynamics and simple vibrational models. Although lattice dynamics reproduce most of the diffuse pattern, protein internal dynamics, which include hinge-bending motions, are needed to explain the short-ranged correlations revealed by Patterson analysis. These insights lay the groundwork for animating crystal structures with biochemically relevant motions.

## Introduction

Conventional structure determination by X-ray crystallography relies on the intense spots recorded in diffraction images, known as Bragg peaks, that represent the average electron density of the unit cell. The average electron density is blurred when atoms are displaced from their average positions, leading to a decay in the Bragg intensities and giving rise to a second signal: a continuous pattern known as diffuse scattering (*1, 2*). Although disorder is routinely modeled in structure refinement of Bragg data as atomic displacement parameters (ADPs) or B-factors (*3*), information about whether groups of atoms move independently or collectively is contained only in the diffuse scattering (Fig. S1). However, the diffuse signal is weak compared to Bragg data and challenging to accurately measure. Diffuse scattering has therefore been largely ignored in macromolecular crystallography, and instead, atomic motions have been inferred solely from Bragg data (*4–6*).

The potential of diffuse scattering as a probe of protein dynamics was envisioned over 30 years ago when Caspar, et al. (*7*) attributed the cloudy diffuse signal from an insulin crystal to liquid-like internal motions. More recently, it has been proposed that diffuse scattering can also disambiguate common structure refinement models that fit collective motions of atoms to ADPs (*8*). Motivated by these key ideas, a number of models of protein motion have been proposed to explain macromolecular diffuse scattering (*2, 9–13*). However, in all cases to-date, agreement between measurement and simulation has been far from compelling (*14–20*), and thus, the promise of diffuse scattering has not yet been realized.

The main bottleneck in the field has been the lack of accurate data. In particular, the diffuse pattern is typically a small oscillation on top of a large background and is therefore easily corrupted by intense Bragg peaks. Thus, it has been common practice to heavily process images either by filtering or masking near-Bragg pixels (*14, 17*). However, this treatment suppresses features that are derived from long-ranged correlations extending beyond the unit cell and may also alter the information contained in the remaining signal. The emerging view is that long-ranged correlations must be considered (*2, 19, 21*), but despite the advent of pixel array detectors that are newly enabling (*22, 23*), diffuse scattering data capable of testing such models have not been reported.

To understand the fundamental origins of diffuse scattering from protein crystals, we analyzed the total scattering from the triclinic form of hen lysozyme (Fig. 1A) collected at ambient temperature using a photon-counting pixel array detector (Fig. S2A). The triclinic crystals (*24*) feature low mosaicity and importantly, one protein molecule per unit cell, ensuring that features between the Bragg peaks are fully resolved. By combining high quality experimental data with new processing methods, we were able to construct a highly detailed map of diffuse scattering without filtering the images. This map reveals, for the first time, a surprisingly large contribution of long-ranged correlated motions across multiple unit cells, while also enabling detection of protein motions in a manner that is consistent with both Bragg diffraction and diffuse scattering.

**Fig. 1:**
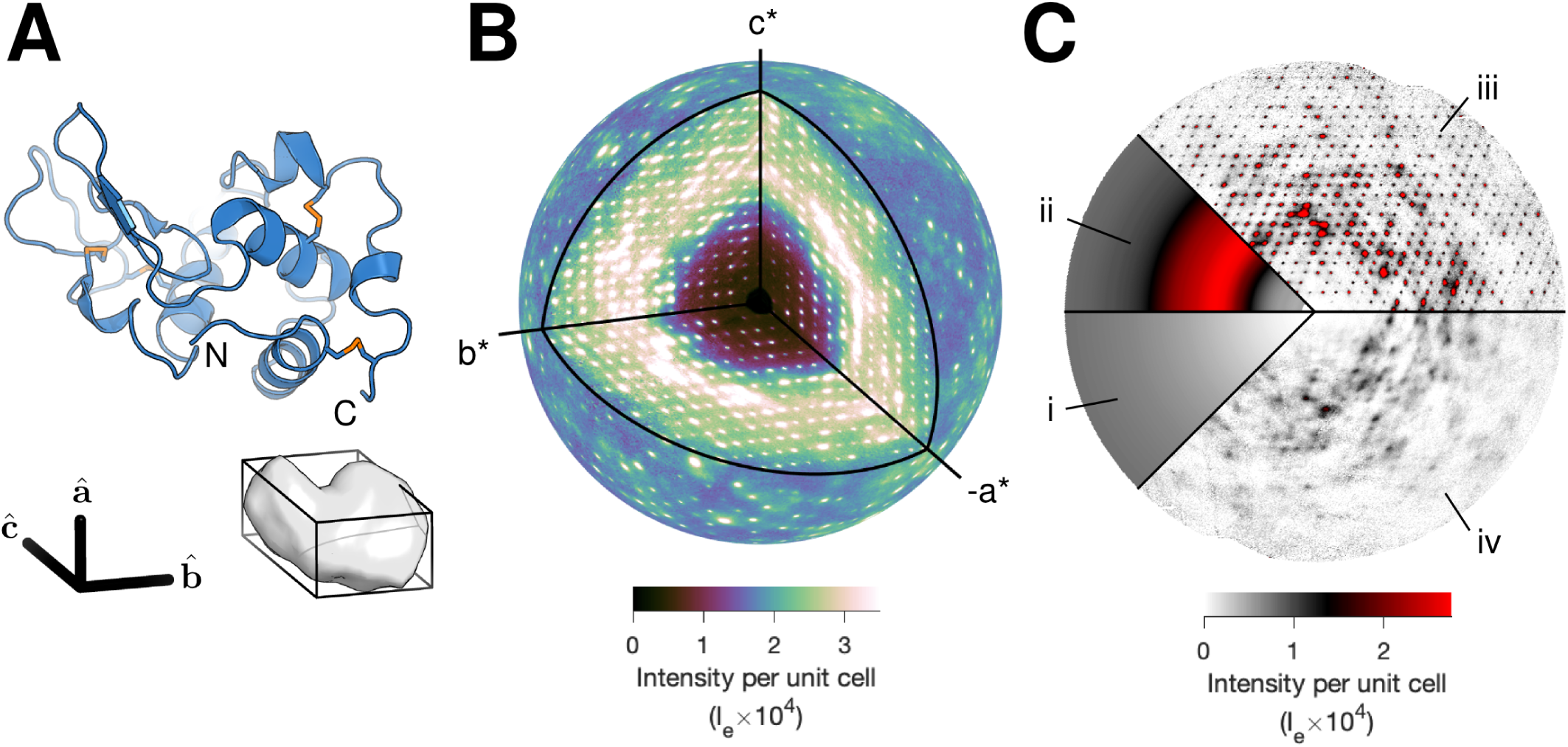
Diffuse scattering map of triclinic lysozyme with intensities on an absolute scale of electron units (*I*_*e*_). (A) Ribbon diagram of lysozyme (top) and the triclinic unit cell containing one protein (bottom). (B) A highly detailed three-dimensional map of diffuse scattering was obtained. The outer sphere is drawn at 2 Å resolution. (C) The total scattering is made up of three components: (i) inelastic Compton scattering, (ii) a broad isotropic ring that dominates the diffuse signal, and (iii, iv) variational features in the diffuse scattering. Intense halos are visible in the layers containing Bragg peaks (*l* = 0 plane, iii). Cloudy scattering is best visualized in the planes mid-way between the Bragg peaks (*l* = 1/2 plane, iv).

## Results

### Construction of a three-dimensional reciprocal space map

For accurate measurements of diffuse scattering at room temperature, the main challenges are to avoid contamination by Bragg peaks and background scattering and to achieve high signal-to-noise while avoiding radiation damage. Using well-collimated and monochromatic synchrotron radiation, we measured the angular broadening (apparent mosaicity) of our triclinic lysozyme crystals to be 0.02 to 0.03 degrees, which is as small as could be resolved by the diffraction instrument (*25*). With such low mosaicity, the sharp, Gaussian-shaped Bragg peaks are readily distinguished from the underlying diffuse scattering (Fig. S3A). To take advantage of this low mosaicity, data were collected with fine phi-slicing (0.1 deg). Crystals were held in low-background capillaries (Fig. S2A), and low-dose partial datasets were collected from multiple sample volumes. In total, four crystals yielded 5500 images from 11 different sample volumes (Fig. S2B, Table S1). Using standard crystallography methods, we determined a structure to 1.21 Å (Table S2) that agrees well with a previously reported room-temperature structure (PDB ID *4lzt* (*24*), 0.14 Å r.m.s.d.). Analysis of the structure and Bragg intensities shows that radiation damage effects were minimal (Fig. S4).

A three-dimensional diffuse map (Fig. 1B) was constructed from the same set of images (described in detail in Supplementary Methods). Background scattering varied with spindle angle (Fig. S2A, S5) and was therefore subtracted frame-by-frame (Fig. S2C). Scale factors for each image pixel were calculated from first principles to account for X-ray beam polarization, detector absorption efficiency, solid angle, and attenuation by air. Additionally, we utilized the high data redundancy to correct for other experimental artifacts, including self-absorption of the crystal, changes in illuminated volume, differences in efficiency among the detector chips, and excess scattering from the loop and liquid on the surface of the crystal (Fig. S6). Each of these corrections improved data quality (Fig. S7). The data were accumulated on a fine reciprocal space grid such that the Bragg peaks were entirely contained within the voxels centered on the reciprocal lattice nodes (Fig. S3B). In this grid, the reciprocal lattice vectors **a**\*, **b**\*, and **c**\* are subdivided by 13, 11, and 11, respectively. The map had a maximum resolution of 1.25 Å, and Friedel pairs were averaged, for a total of ∼50 million unique voxels.

To enable rigorous comparison between simulations and experiment, we adapted the integral method of Krogh-Moe (*26, 27*) to place the map on an absolute scale of electron units per unit cell (Supplementary Methods, Fig. S8). By doing so, we are able to subtract the inelastic scattering contribution, which depends only on the atomic inventory and is insensitive to molecular structure (Fig. 1C, i). The final diffuse map thus represents coherent scattering features of interest (Fig. 1B), which depends on structure.

### Phonon-like scattering

The diffuse scattering is dominated by a broad, isotropic scattering ring with a peak at ∼ 3 Å (Fig. 1C, ii) that arises from short-ranged disorder generally attributed to water (*28, 29*). To better visualize the non-isotropic fluctuations, we resampled the full map mid-way between the Bragg peaks and defined the isotropic background as one sigma level below the mean scattering of this map in each resolution bin (Supplementary Methods, Fig. S9). Subtracting this background from the full map reveals clear non-isotropic features, hereafter referred to as “variational” (Fig. 1C, iii-iv). The most striking variational features are the intense halos (Fig. 1C, iii) that appear to co-localize with Bragg peaks at the reciprocal lattice nodes (Fig. 1B), which are significantly asymmetric in certain directions (Fig. S10, left). Overlaid with the halos is a cloudy pattern that is found throughout the map (Fig. 1C, iv), which we estimate accounts for roughly half of the integrated variational intensity in most resolution bins (Fig. S11).

The presence of such clear halo scattering was unexpected as it implies that there are correlations between atoms in different unit cells. In protein crystallography, an outstanding question has been whether such correlations are dynamic in nature, and specifically, due to lattice vibrations (*7, 9, 15, 21, 28, 30*). The scattering intensity of a phonon (vibrational mode) is proportional to the mean squared amplitude of vibration and peaks at certain points in reciprocal space. In particular, a phonon with wavevector **k** makes the greatest contribution when the scattering vector **q** (with magnitude |**q**| = 2*π/d*) is parallel to the phonon polarization and displaced from the nearest Bragg peak at **q**_0_ such that **q** − **q**_0_ = *±***k** (*31*). The scattering of the so-called acoustic phonons, which are thermally excited at room temperature, is proportional to *v s*^−2^ |**k**|^−2^, where *v*_*s*_ is the speed of sound. Thus, at the Bragg peak locations, acoustic phonon scattering is expected to produce halos with a characteristic |**q** − **q**_0_|^−2^ decay in intensity in any given direction.

With our finely sampled diffuse map, the halo scattering can be inspected directly. We selected three symmetric and intense halos and plotted their intensities along the three reciprocal axes on a double-log scale, where a power law is a straight line (Fig. 2A, left). Both the power-law behavior and the characteristic exponent are fully consistent with acoustic phonon scattering. Furthermore, the fact that the plot remains linear as **q** approaches **q**_0_ implies that the lattice vibrations are coherent over at least 2*π/*|**k**_min_| ∼ 300 Å or ∼ 10 unit cells. The characteristic exponent of approximately −2 is also found for other intense halos throughout the map (Fig. 2A, right). These results are highly suggestive of vibrational lattice dynamics.

**Fig. 2:**
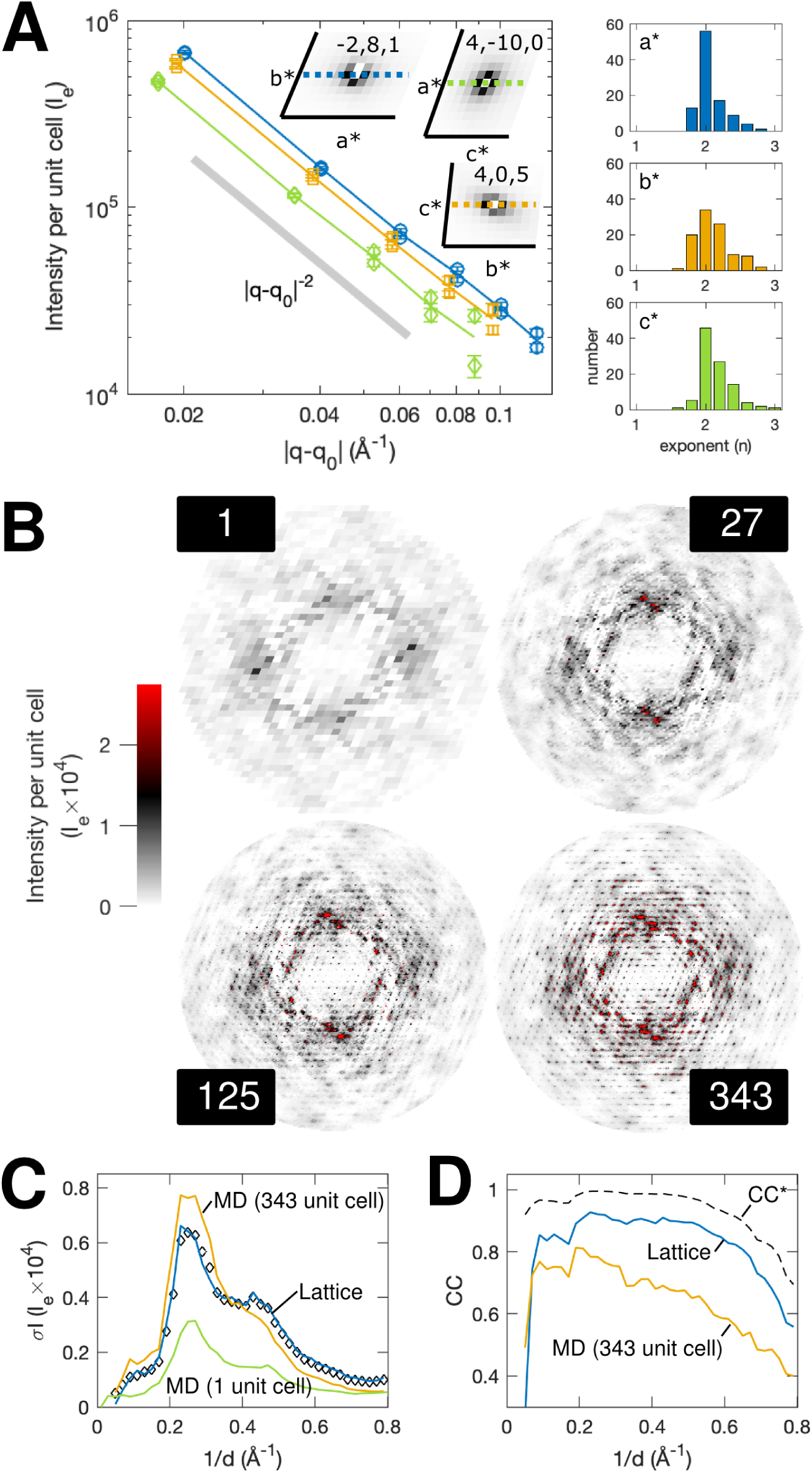
Evidence for long-range correlations in experimental maps and molecular dynamics (MD) simulations. (A) Throughout the diffuse map, intense halo scattering is observed around Bragg reflections. Halo profiles centered on three Bragg reflections (**q**_0_) show a power-law decay with an exponent close to −2 (gray line) along the directions (**q** − **q**_0_): **a**\* (blue), **b**\* (orange), and **c**\* (green). Histograms of the best-fit exponent along **a**\*, **b**\* and **c**\* (top to bottom) for the 100 most-intense halos between 2 and 10 Å resolution also show that −2 is the most frequent value. (B) Halo features appear in simulated scattering from supercell MD as the simulation size is increased from 1 to 343 (7×7×7) unit cells. Each panel shows the variational component in the *l* = 0 plane. (C) Although increasing the supercell size improves agreement (green to orange), MD does not reproduce experiment on an absolute scale (black diamonds), as judged by the standard deviation profile of the diffuse intensity. In contrast, much better agreement is obtained with the lattice model described in Fig. 3 (blue). (D) MD displays a worse correlation (CC) with experiment (orange) compared to the lattice model (blue). The dashed line represents theoretical limit of the experimental data, CC*.

### All-atom molecular dynamics simulations

Although all-atom MD simulations have previously been used to investigate the contribution of protein dynamics to diffuse scattering (*12, 16, 29, 32–34*), the effect of long-ranged correlations due to lattice disorder has not been examined. We thus performed all-atom MD simulations of triclinic lysozyme crystals as a function of supercell size (Supplementary Methods). Experimentally determined coordinates were used to define and initialize an array of proteins comprising the supercell, and periodic boundary conditions were imposed to remove edge effects. Supercells composed of 1, 27 (3×3×3), 125 (5×5×5), and 343 (7×7×7) unit cells were simulated for 5, 5, 2, and 1 *µ*s, respectively. Guinier’s equation (*35*) was used to calculate the diffuse intensity per unit cell from the simulation trajectory (Supplementary Methods). Because the boundary conditions are periodic, the diffuse scattering was sampled at integer subdivisions of the reciprocal lattice (i.e. the number of unit cells in each direction).

In the 1 unit-cell simulation (Fig. 2B), cloudy variational features are observed in rough qualitative agreement with the experiment (Fig. 1C, iii-iv), suggesting that local protein and solvent dynamics contribute to the observed diffuse scattering. Unlike simpler models that do not include liquid correlations in the bulk solvent, MD provides a prediction for the isotropic component (Fig. S9C). The overall correlation of the isotropic component is 0.9965 between 25 and 1.25 Å resolution, and the magnitude is also similar (Fig. S8A,C). However, halos are absent, consistent with the lack of intermolecular disorder enforced by a 1 unit-cell simulation. As the size of the supercell is increased, the diffuse scattering pattern evolves in a complex manner with the halos becoming increasingly apparent (Fig. 2B), confirming that they depend on intermolecular correlations and lattice degrees of freedom. In the 343 unit-cell simulation, the r.m.s. displacement of each chain about its center of mass was 0.20 to 0.22 Å in each direction. Although this may seem to be a small motion, the intense halo signal is derived from the collective motions of many proteins.

In the 343 unit-cell simulation (Fig. 2B), the simulated scattering contains both cloudy and halo features similar to those observed experimentally. To make a quantitative comparison, we interpolated the experimental map on the simulation grid (7×7×7) and computed the Pearson correlation coefficient (CC) between the two in thin shells of constant resolution (Fig. 2D, orange). Although we obtain a reasonable CC of ∼ 0.7 up to 2 Å resolution, the CC decreases at higher resolution. Moreover, there is a significant gap between CC (Fig. 2D, orange) and CC* (Fig. 2D, black dashed), which estimates the maximum CC a model can achieve, given the precision of the data (*36*). This discrepancy indicates that model-data agreement is not limited by noise and instead points to shortcomings of the crystal model, including the current MD force fields. In particular, the accuracy of MD for diffuse scattering appears to be limited by errors in the average electron density (Fig. S12). To gain insight into the underlying physics of the variational scattering features, we thus sought simpler dynamical models that can be refined to fit both the Bragg and diffuse data.

### Lattice dynamics refined against diffuse scattering

Given the evidence for acoustic phonon scattering, we investigated whether vibrational models can capture the observed halo shapes and intensities. We developed a lattice dynamics model where each protein is able to move as a rigid body that is connected to neighboring molecules via spring-like interactions (detailed in Supplementary Methods). The proteins were arranged in a 13×11×11 supercell to match the sampling of the experimental map (Fig. 3A). Residues of neighboring molecules that form lattice contacts were linked by a pair potential between alpha carbons (Fig. 3A, magenta lines), reflecting a restoring force that depends on the relative displacements of the two end-points. For generality, we allowed each pair potential to be a linear combination of two types of springs: Gaussian and directional. Gaussian springs (*37*) have a restoring force that is independent of the direction of the displacement relative to the spring, and directional springs (*38*) have a restoring force only along the vector between the end points.

**Fig. 3:**
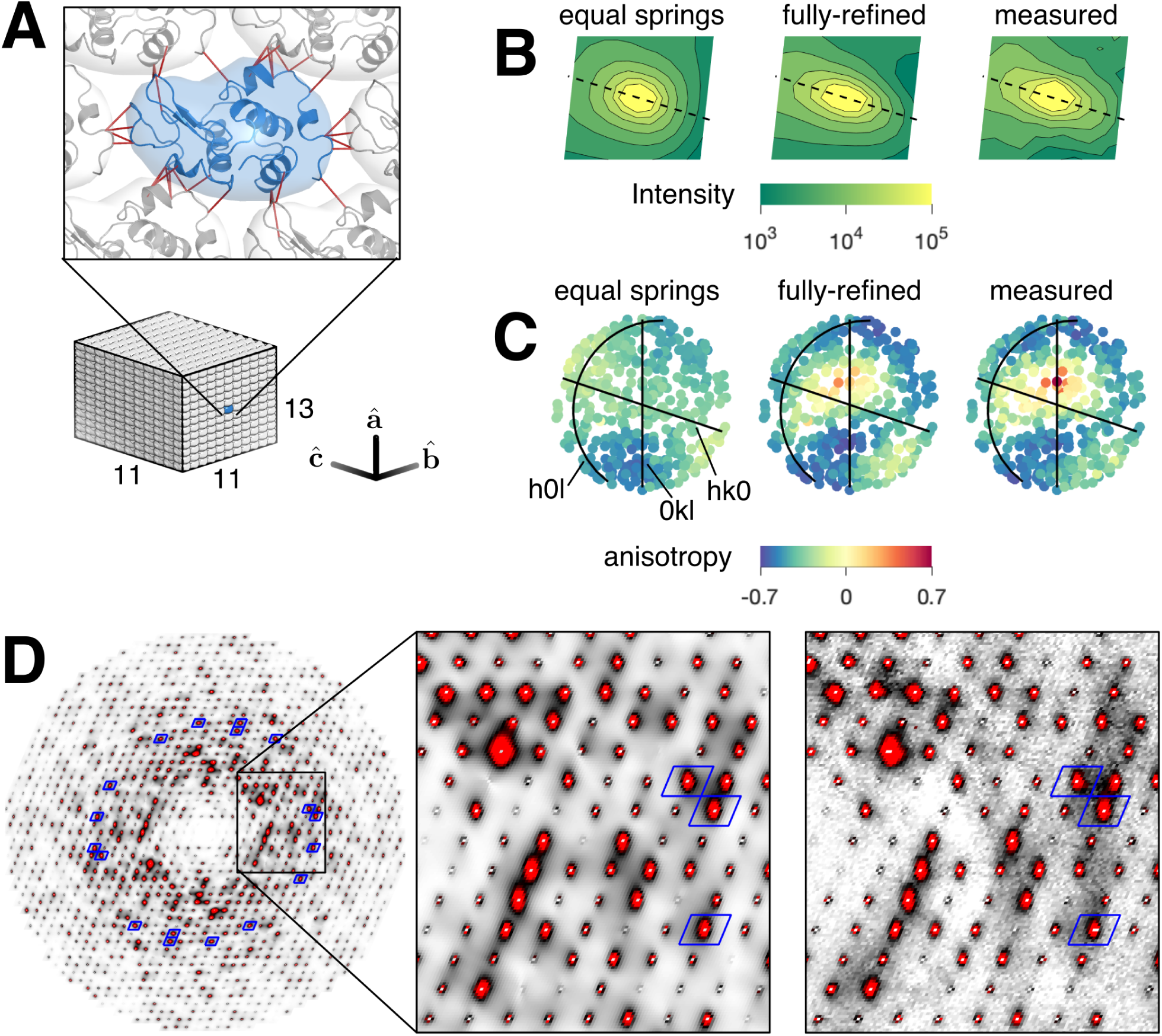
Lattice dynamics model refined to diffuse scattering. (A) A lattice dynamics model was constructed with rigid protein units arranged in a 13×11×11 supercell with a linear combination of Gaussian and directional springs connecting the C_*α*_ atoms of residues involved in lattice contacts (dark red lines). Spring constants were refined to fit the variational scattering around 400 intense Bragg peaks between 2 and 2.5 Å resolution. (B) Comparison of predicted and measured halo intensity around the (1,2,13) Bragg reflection in the *h* = 0 plane. The plane perpendicular to the scattering vector is indicated by a dashed line. The model with equal Gaussian springs does not reproduce this shape as well as the fully-refined model. (C) The shape anisotropy of each of the 400 halos used for model fitting was quantified and mapped as an equal-area projection of the hemisphere centered on **b**\*. Full refinement of the spring constants was needed to reproduce the pattern of halo anisotropy seen in experiment. (D) The simulated one-phonon scattering for the fully-refined lattice model (left) is compared with the measured variational scattering (right) in the *l* = 0 plane. The intensity scale is the same as Fig. 1C. Blue boxes surround halos that were included in the fit.

The model was refined against a set of 400 intense halos between 2 and 2.5 Å resolution, consisting of a total of ∼600,000 voxels. As there are halos associated with all 30,108 unique Bragg reflections, these 400 represent a small subset (1.3%). The spring constants were initially restrained to be all Gaussian and equal, and restraints were relaxed during subsequent stages of refinement. For a given set of springs, the equations of motion were solved by the Born/Von-Karman method (*31, 39, 40*), and the diffuse scattering was calculated using the one-phonon approximation (detailed in Supplementary Methods). At each refinement stage, we monitored the overall *χ*^2^ value between the experimental and simulated scattering (Fig. S13A), as well as the ability of the model to reproduce the halo shape (Fig. 3B). To monitor agreement with halo anisotropy, we fit each of the halos to a function of the form *I* = [(**q** − **q**_0_)**G**(**q** − **q**_0_)]^−1^, where **G** is a 3×3 positive definite matrix, and defined an anisotropy parameter, *a* = **G**_⊥_*/***G**_‖_ − 1, where **G**_‖_ is the component of **G** parallel to **q**_0_, and **G**_⊥_ is the average of the perpendicular components. The fully-parameterized model was necessary to reproduce the pattern of halo anisotropy (Fig. 3C, Fig. S13B).

After refining the lattice dynamics model using the working set of 400 halos (Fig. 3D, blue boxes), we simulated the complete diffuse scattering map over the full resolution range. Remarkably, the simulation reproduces many of the variational scattering features observed in experiment (Fig. 3D, right). Anisotropic halo shapes are reproduced even in regions of the map that were not used to refine the model (Fig. 3D, regions outside of blue boxes). Streaks in the pattern are also reproduced and can be attributed to a modulation of the halos by the molecular transform (Fig. S10). Moreover, we find that the halos do not decay to zero mid-way between the Bragg peaks as previously expected (*9*), giving rise to a cloudy pattern that resembles the cloudy variational scattering in the data (Fig. 3D, right). The standard deviations of intensity have very similar profiles and absolute magnitudes (Fig. 2C, blue solid and diamonds), suggesting that the lattice dynamics make the most significant contribution to the scattering variations. This conclusion is supported by the much smaller variations seen in the 1 unit-cell MD simulation (Fig. 2C, purple), where lattice disorder is absent by construction.

As before, the agreement between the experimental and simulated maps was assessed with CC and CC*. For the lattice dynamics model, the CC is excellent in regions of high signal-to-noise (CC ∼ 0.9 between 2 and 5 Å resolution) (Fig. S14, solid) and only limited by the experimental precision at higher resolution (Fig. S14, dashed). To improve the signal-to-noise, the maps were interpolated on a 7×7×7 grid (Fig. 2D, blue), enabling direct comparison with the 343 unit-cell MD simulation (Fig. 2D, orange). Strikingly, the lattice dynamics model clearly outperforms all-atom MD (Fig. 2C-D).

The lattice model can be further assessed against existing biophysical data. Our model predicts that sound waves should propagate through the crystal. Based on the calculated dispersion relations of the acoustic vibrational modes (Fig. S15), we obtain longitudinal sound velocities of 1.0-1.3 km/s and corresponding transverse velocities that are slower by a factor of 1.3-2.1 depending on the propagation direction (Table S3). Although few measurements of sound propagation have been made in protein crystals, longitudinal velocities have generally been reported to be ∼2 km/s (*41–43*), and transverse velocities are estimated to be 2-3 times slower (*41, 44*). Thus, our interpretation that the halo scattering arises from dynamic, rather than static, disorder appears physically reasonable.

### Contribution of lattice dynamics to atomic motion

As described earlier, the amount of apparent motion for each atom can be quantified from Bragg data by refining individual ADPs, the 6 components needed to describe a 3-dimensional Gaussian probability distribution. Our data quality was sufficient to refine full anisotropic ADPs for every non-H atom. To determine the extent to which lattice dynamics contribute to atomic motion, corresponding ADPs were calculated directly from the refined lattice model (Supplementary Methods, Table S4). In Fig. 4A, the full ADPs of the backbone atoms are reduced to a single isotropic B-factor per residue to facilitate visual comparison. Overall, the backbone B-factors for the lattice model (5.2 Å^2^ on average) fall below those of experiment (9.4 Å^2^ on average). The B-factors from the lattice model show small variations, which can be attributed to rigid-body rotational motion with an r.m.s. amplitude of 0.8° (Table S4). However, the B-factor variations in the data are much more pronounced (Fig. 4A), particularly for side-chains (Fig. S16A). These residual B-factors imply the existence of internal dynamics, in other words, that atoms within the protein undergo collective motions.

**Fig. 4:**
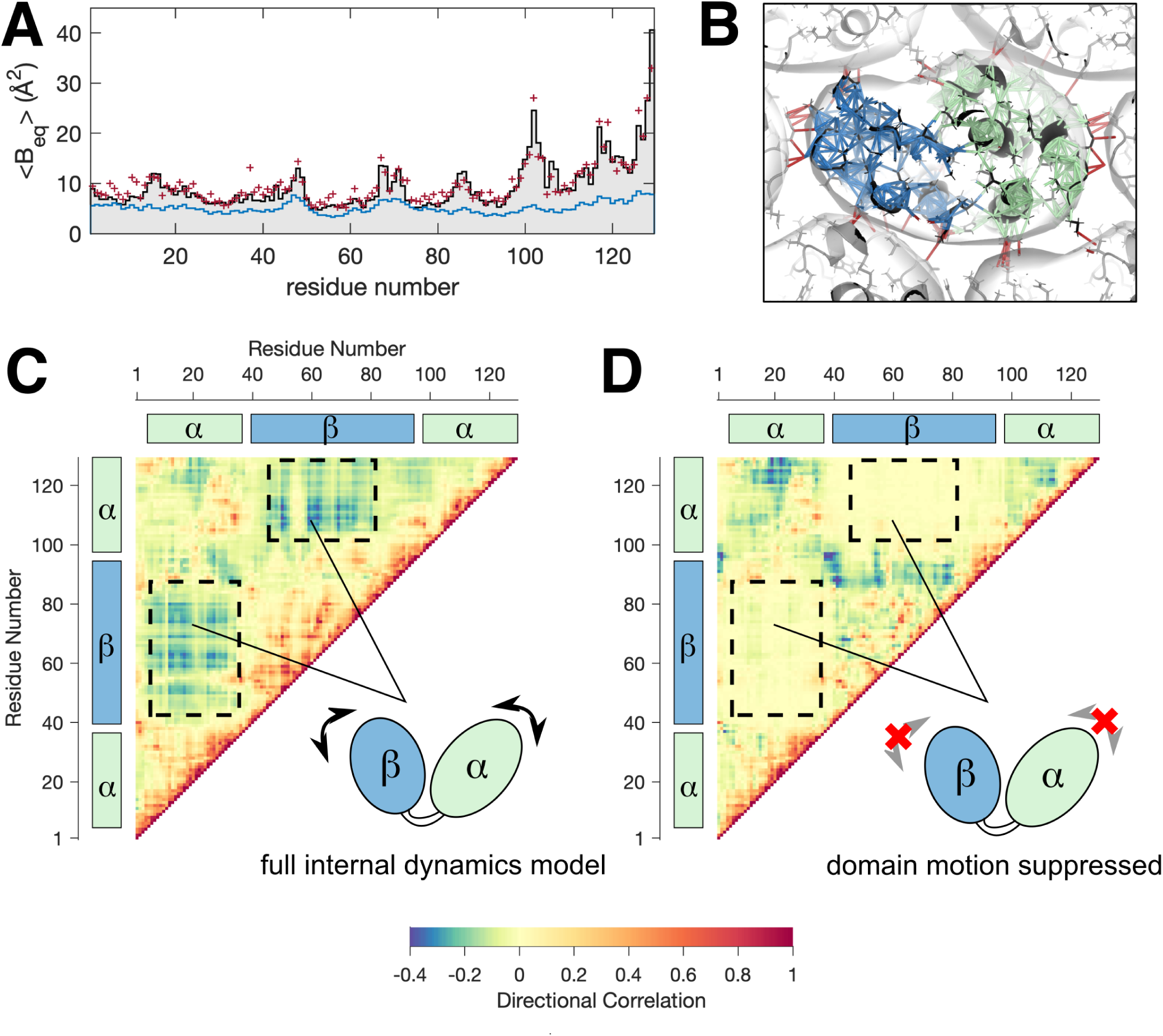
Models of collective internal motions in lysozyme refined to Bragg data. (A) Apparent atomic motions can be evaluated by comparing the anisotropic displacement parameters (ADPs) obtained experimentally from Bragg data with those calculated from models. To facilitate visual comparison, the ADPs over backbone atoms are averaged to produce a single isotropic B-factor per residue. The lattice dynamics model in Fig. 3 (blue curve) underestimates the experimental B-factors (gray bars), but a good fit is obtained by combining lattice dynamics with the internal dynamics described in panels B-C (dark red symbols). (B) The model for internal dynamics was constructed using an elastic network with rigid residues. Both intermolecular (dark red lines) and intramolecular contacts (blue and green lines, corresponding to the *α* and *β* domains, respectively) were modeled as springs, and the spring constants were refined to fit the residual ADPs, i.e. the experimental ADPs that are unaccounted for by lattice dynamics. (C) In the full internal dynamics model, the C_*α*_ atoms in the *α* and *β* domains show negative directional correlations (dashed boxes), indicating that their motions are anti-correlated and consistent with hinge-bending. (D) The two domains have no correlations when their motions are suppressed in the model refinement.

### Protein dynamics refined against Bragg data

The collective motions of lysozyme have been a topic of long-standing biophysical interest since hinge-bending motions between the two domains (Fig. 4B, blue and green) were first proposed as a mechanism for substrate binding and release (*45, 46*). To investigate the presence of such collective motions, we developed an elastic network model, in which each protein residue moves as a rigid body, and all non-H atoms within 4 Å are coupled with directional springs (Supplementary Methods). As with the lattice model, the crystal environment was modeled with intermolecular springs and periodic boundary conditions, and the dynamics were calculated using the Born/Von-Karman method. In order to model only the internal protein dynamics, the Hessian matrix describing the restoring forces was modified to suppress rigid-body motion of the entire protein. The model was parametrized with one coupling constant per residue (i.e. 129 free parameters total) so that springs joining a residue pair were assigned a spring constant equal to the geometric mean of the coupling constants (Supplementary Methods). The parameters were then refined by minimizing the least-squares difference between all components of the calculated (lattice + internal) and experimental ADPs derived from Bragg data.

The refined model is able to reproduce the pattern of B-factors obtained experimentally (Figs. 4A and S16A,B). To assess the importance of hinge-bending in the model, we examined the covariance matrices **C**_*ij*_ for all alpha carbon pairs and calculated a “directional correlation”, which is the component of **C**_*ij*_ along the inter-atomic vector normalized by the r.m.s. displacements of the two atoms (Supplementary Methods). By this measure (Fig. 4C), the two domains are significantly anti-correlated as expected for hinge-bending motion.

### Contribution of protein dynamics to diffuse scattering

Lattice dynamics account for the bulk of the variational diffuse scattering, as evaluated by CC and standard deviation (Fig. 2C-D). However, these statistics emphasize the most intense features in the signal, which in this case are the halos. To assess the more subtle contributions of internal protein motions, correlations in the signal should be separated based on length-scale. We thus calculated the diffuse Patterson (also known as 3D-ΔPDF), which is the Fourier transform of the diffuse scattering. The diffuse Patterson map represents the mean autocorrelation of the difference electron density, Δ*ρ* = *ρ*−⟨*ρ*⟩, such that a vector from the origin of the map corresponds to a vector between two points in the crystal. Thus, the central part of the diffuse Patterson is affected only by those correlations that are short-ranged.

At large distances, the experimental diffuse Patterson displays peaks at the lattice nodes as expected (Fig. 5A, left), whereas continuous features are most intense at short distances (Fig. 5A, right). To determine whether lattice dynamics alone can account for the short-ranged correlations, the diffuse Patterson was calculated directly from the refined lattice model (Supplementary Methods). Although the simulated and experimental maps share similar features (Fig. 5A-B), the amplitudes of the fluctuations are clearly underestimated for distances shorter than ∼ 10 Å (Fig. 5F, blue curve vs. diamonds).

**Fig. 5:**
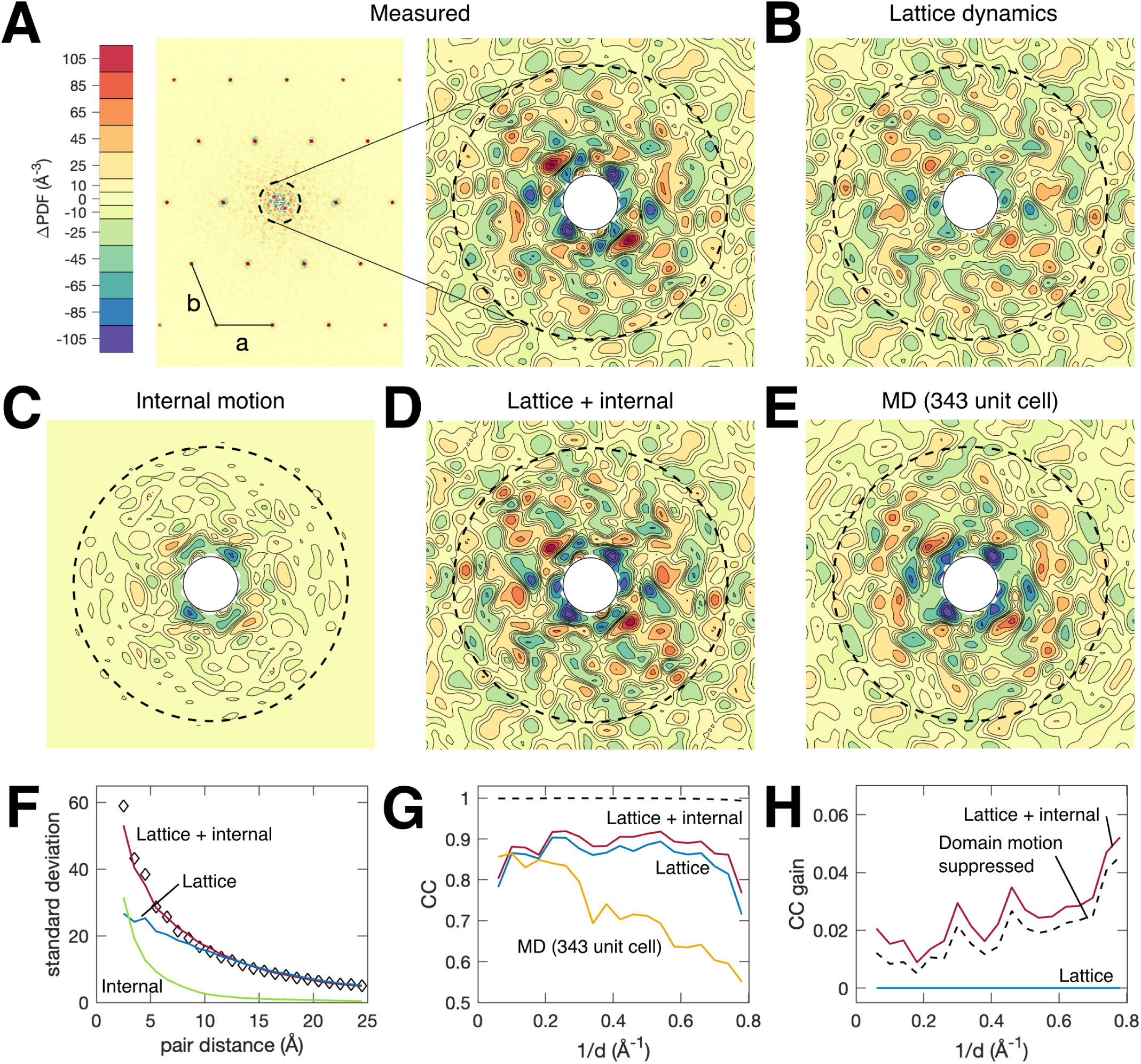
Detection of internal motions by diffuse Patterson analysis. (A) The diffuse Patterson map represents the autocorrelation of the difference electron density and is a function of the vector **r** between points in the crystal (dashed circle corresponds to |**r**| = 10 Å). The experimental diffuse Patterson in the **a**-**b** plane contains peaks at the lattice nodes (left) and continuous fluctuations that are most intense near the origin (right). (B-E) Diffuse Patterson maps simulated from models. The lattice model underestimates the fluctuations at short length scales, but addition of full internal dynamics reproduces the experimental pattern. (F) The standard deviation of the diffuse Patterson maps in spherical shells of constant pair-distance for the experimental map (black diamonds), lattice model (blue), internal model (green), and the combined model (dark red). (G) The reciprocal space correlation coefficient (CC) between experiment and simulation in shells of constant resolution within the central part of the Patterson map (2 < |**r**| < 25 Å). The colors are the same as in panel F. The 343-unit cell MD simulation is in orange. The dashed line shows CC*. (H) Gain in CC relative to the lattice dynamics model alone (blue) for the combined model (dark red) and the model in which domain motion was suppressed (black dashed line).

In contrast, the diffuse Patterson calculated from the refined internal motion model shows prominent fluctuations for pair distances less than ∼ 10 Å but very little outside this range (Fig. 5C and Fig. 5F, green). Assuming that the protein internal motions are independent of lattice motions, the diffuse Patterson maps can be added (Fig. 5D). The combined model displays remarkable agreement with the experimental map and reproduces the characteristic decay of fluctuation amplitude almost exactly (Fig. 5F, dark red curve vs. diamonds). To assess the agreement more quantitatively, the CC profile was calculated in reciprocal space (a Fourier transform of the 2 < *r* < 25 Å region). The combined model (Fig. 5G-H, dark red) displays a significant gain in CC over the lattice model alone (Fig. 5G-H, blue). The level of model-data agreement that we obtain is excellent (Fig. 5G, dark red), especially when compared to the all-atom MD simulation (Fig. 5E and Fig. 5G, orange) as well as all previously reported studies (*14, 15, 17–20, 33*). However, it is still less than the theoretical maximum of our diffuse Patterson map (Fig. 5G, dotted). CC* is ∼1 for most resolution bins as truncating the Patterson function is equivalent to band-pass filtering the intensities, and it greatly increases signal-to-noise. Thus, there is room for improvement if better models can be devised.

The question of model quality has consequence to protein crystallography, where it is common practice to fit models of collective motion to the B-factors, since this often increases the data-to-parameter ratio. Diffuse scattering has been proposed as a means of critically evaluating these models (*8*). To explore this idea, we repeated refinement of the internal elastic network model with domain motions selectively suppressed. Although the internal dynamics are significantly different in this model (Fig. 4D), they also reproduce the experimentally derived B-factors (Fig. S16C). This result underscores the challenges of distinguishing differing models of protein motion from Bragg data alone. However, fluctuations in the diffuse Patterson decay more rapidly with domain motions suppressed (Fig. S17), leading to a subtle but systematically worse CC, particularly at high resolution (Fig. 5H, dashed).

## Conclusions

By studying the total X-ray scattering from triclinic lysozyme crystals both experimentally and theoretically, we were able to obtain fundamental insight into the collective motions that produce macromolecular diffuse scattering. Simple vibrational models of the lattice and internal dynamics were developed that explain the electron density correlations spanning two orders of magnitude in length-scale. Vibrations of the entire protein in the lattice account for the shapes and magnitudes of the diffuse halo features and about half of the backbone ADPs, while internal motions of the protein make up the remainder. The collective nature of these internal motions was investigated by diffuse Patterson analysis, which separates correlations based on the inter-atomic vector. Remarkably, we found that two models that fit the ADPs equally well could be distinguished by their agreement to the experimental diffuse Patterson, experimentally demonstrating a key application of diffuse scattering proposed a decade ago (*8*). Finally, although the MD was limited in its ability to reproduce the variational diffuse scattering, our results demonstrate that this signal provides an excellent experimental benchmark for improving simulations in the future.

For over 30 years, the ultimate goal of diffuse scattering studies has been to capture internal protein motions from crystallographic data. The success of previous efforts has been limited primarily by data quality as well as the assumption that the variational scattering is largely due to internal motion. In fact, lattice disorder contributes significantly, further underscoring the need for detailed, high quality data and realistic models. Despite the added challenges, we have also shown that by accounting for lattice dynamics, the remaining diffuse signal indeed contains information about internal motion and can be used to differentiate alternate models. With the initial goal of the diffuse scattering field realized, the next grand challenge of refining structural models that are consistent with the total scattering now appears within reach.

## Acknowledgments

We thank staff at the Cornell High Energy Synchrotron Source (CHESS) and MacCHESS beamline F1 for supporting diffraction data collection, W.C. Thomas for assistance with data collection, and S.M. Gruner, W.C. Thomas, M.B. Watkins, B.R. Crane, and A.S. Byer for critical reading of the manuscript.

## Funding

CHESS is supported by NSF Grant DMR-1332208, and the MacCHESS facility is supported by NIH/NIGMS Grant GM-103485. This work was supported by NIH Grants GM117757 (to S.P.M.), GM100008 (to N.A.), GM124847 (to N.A.), and GM122086 (to D.A.C.) and by start-up funds from Princeton University and Cornell University (to N.A.).

## Author contributions

S.P.M. performed crystallization, data collection, and structure determination with assistance from N.A. D.A.C. performed and analyzed the MD simulations. S.P.M. developed methods for processing and analyzing diffuse scattering data and performed vibrational simulations. The manuscript was written by S.P.M. and N.A. and edited by all authors. N.A. conceived of the experiments and coordinated the research.

## Competing interests

The authors declare no competing interests.

## Data and materials availability

The atomic coordinates and structure factors have been deposited in the Protein Data Bank under accession code *6o2h*. Diffraction images have been deposited in the SBGrid Data Bank. All other data are available in the main text or the supplementary materials.

## Supplementary Materials

Materials and Methods

Supplementary Text

Fig. S1 – S17

Table S1 – S4

