## Supplementary Materials for "Diffuse X-ray Scattering from Correlated Motions in a Protein Crystal"

**This PDF file includes:**

- Materials and Methods
- Supplementary Text
- Figs. S1 to S17
- Tables S1 to S4

### Materials and Methods

#### Crystallization

Triclinic crystals of lysozyme were obtained by the micro-batch method with temperature-cycling to select this crystal form based on its phase diagram (*1*). Lyophilized hen egg white lysozyme (Hampton Research) was dissolved in 20 mM sodium acetate (NaOAc) pH 4.6 at a stock concentration of 100 mg/mL, passed through a 0.2 $\mu$m filter, and used without further purification. All other reagents were purchased from Sigma, unless noted. Triclinic crystals were grown using a microbatch-under-oil technique with 8 $\mu$L drops containing 5-15 mg/mL protein, 224-300 mM $NaNO_3$, and 50 mM NaOAc pH 4.5 that were covered with 20 $\mu$L paraffin oil. Crystallization trays were set up at room temperature, moved to 4 °C for 8-12 hours, and then returned to room temperature. Both triclinic and monoclinic crystals nucleate during the 4 °C incubation, however after returning to room temperature the triclinic crystals grow at the expense of the monoclinic form (*1*).

#### Data collection

X-ray data were collected using the macromolecular crystallography beamline F1 at the Cornell High Energy Synchrotron Source (CHESS), which provided a 12.693 keV X-ray beam collimated to 0.1 mm diameter. Room-temperature data collection was performed using the plastic capillary sheathing method (*2*). Crystals were harvested using low-scatter kapton loops (MicroLoops, MiTeGen), taking care to minimize the amount of solvent surrounding the crystal, and placed within 2 mm diameter, 25 $\mu$m wall poly(ethylene terephthalate) capillaries (MicroRT, MiTeGen) with 10 $\mu$L reservoir solution in the tip. During X-ray exposure, images were recorded every 0.1° using a pixel-array detector (Pilatus3 6M, Dectris) while rotating the sample at $1^\circ\,s^{-1}$. A dose rate of 1.3 kGy $s^{-1}$ was estimated assuming a flux of $2.5 \times 10^{10}$ photons $s^{-1}$ and a mass energy-absorption coefficient (*3*) of $\mu_{en}/\rho = 2.0\,cm^2 g^{-1}$. After 50 s of exposure (∼65 kGy), the sample was refreshed by translating to a new spot or replacing the crystal. A background dataset was collected for each crystal by translating the sample out of the beam along the spindle axis and collecting 1 s exposures while rotating at $1^\circ\,s^{-1}$.

#### Structure determination from Bragg data

Bragg data were integrated using xds (*4*), with geometric parameters refined at 2° increments. The fitted peak profiles and mosaicity per frame were examined to verify that the crystal had not slipped or cracked. The best wedges were then scaled and merged using aimless (*5*) (Table S1). Model building and refinement were carried out using the ccp4 suite of programs (*6, 7*). The initial model was prepared from PDB ID *4lzt* (*8*), using the most probable (highest occupancy) protein atom coordinates only. Structure refinement was carried out using alternate runs of REFMAC5 (*9*) and manual modeling in coot (*10*), with atomic displacement parameters included in the final rounds. Alternate conformers were modeled when justified by the electron density and stereochemistry. The atomic coordinates and structure factors have been deposited in the Protein Data Bank under accession code *6o2h*. Data collection and refinement statistics are shown in Table S2.

#### Processing of diffuse scattering data

Reciprocal space maps were generated in Matlab (The Mathworks) as described in the following sections. Briefly, an integration mask was first produced to separate rapidly varying features (including Bragg peaks) from continuously varying features (including diffuse scattering) in three-dimensional reciprocal space. Following per-pixel image corrections, the integration mask was used to generate a coarse continuous scattering map and a Bragg map. Using the coarse continuous scattering map, a scaling model was refined to globally minimize the discrepancy between redundant observations. Bragg intensities were corrected and brought into agreement with the values used for structure determination in the previous section. The intensities were then placed on an absolute scale. Finally, continuous scattering intensities were accumulated on a fine grid to produce the final diffuse map. The scaling corrections from the previous step were applied during integration, and redundant observations were merged without further scaling. A detailed description of each operation is below.

**Construction of integration mask.** As the Bragg intensities and continuous scattering require different corrections, a sensitive moving-window filter was first used to detect and mask out rapidly varying features. The filter algorithm compared the observed count distribution to that expected from Poisson statistics, as described below. Briefly, a voxel was masked out if its exclusion made the neighborhoods to which it contributes more Poisson-like according to the Kullback-Leibler (KL) divergence of the observed and ideal distributions. The unmasked voxels then describe a function that varies smoothly on the scale of the reciprocal space grid.

X-ray images were processed in 2° wedges. Each pixel was mapped onto a provisional reciprocal space grid, where the reciprocal unit cell was subdivided by a factor of 5 in each direction. For each voxel, a histogram of counts per pixel was accumulated. Using these count histograms, the filtering algorithm proceeded as follows. The neighborhood (filter window) was defined as the set of voxels within a Euclidian distance of $\leq 2$ grid units from the central voxel. For each neighborhood, the weighted median count rate $r_{\mathrm{median}}$ was found, as well as the KL divergence of the total count histogram from the expected Poisson distribution with rate $r_{\mathrm{median}}$. A voxel was masked if its exclusion reduced the sum of KL divergences for all neighborhoods. First, the voxels were ranked by this change in KL divergences, $\Delta_{\mathrm{KL}}$, in ascending order (worst offenders first). Then, voxels were masked progressively, and the $\Delta_{\mathrm{KL}}$ values of neighboring voxels were updated without re-sorting. The algorithm halted after encountering a voxel with $\Delta_{\mathrm{KL}} \geq 0$ . In the resulting integration mask, the unmasked voxel grid represented the continuous scattering, whereas the masked voxel grid consisted mainly of Bragg peaks.

**Integration and scaling.** Per-pixel image corrections were applied prior to integration. Scale factors for each image pixel were calculated from first principles to account for X-ray beam polarization, detector absorption efficiency, solid angle, and attenuation by air (see Appendix 2 of the Supporting Text). The background count rate for each pixel was estimated from an exposure where the crystal was translated out of the beam along the spindle axis (Figure S2C).

Using the mask generated in the previous step, the unmasked and masked voxels were then integrated separately in 2° wedges. To generate a coarse map of continuous scattering, the unmasked voxel grid was reduced to one sample per reciprocal lattice node. In addition, observations of the same voxel in adjacent wedges were combined. Then, the geometric and background corrections were applied to the photon counts to generate a map of $I_{\mathrm{meas}}$ for the continuous scattering (Equation 52 in Supporting Text). The masked voxels containing Bragg peaks were integrated in a similar manner, except that the local diffuse background was subtracted and the Lorentz correction was applied (Equation 54). For the background, the value of $I_{\mathrm{meas}}$ for the coarse continuous scattering map was used. The Bragg intensities were further filtered to remove partial observations. The total reciprocal space volume sampled by the detector during integration (the accumulation of Equation 48 over contributing pixels) was compared with the actual volume of the masked voxels. The peak was considered to be fully recorded if the volumes agreed within 5%. This rejects a large fraction of the recorded Bragg peaks, however they are later replaced using more precise integration methods, described below.

Using the coarse continuous scattering map, a scaling model was refined in order to minimize the discrepancy of redundant observations and correct for experimental artifacts. In this case, redundancy comes from Friedel symmetry and the fact that different wedges of data overlapped in reciprocal space. The scaling model related the expected intensity of an observation $i$ to the merged intensity $I_{\mathrm{merge}}$, in terms of four correction factors, as follows:

$$I_{\mathrm{pred}}(i) = a(x_i, y_i, \phi_i) d(p_i) \left[ b(\phi_i) I_{\mathrm{merge}}(\mathbf{h}_i) + c(s_i, \phi_i) \right], \tag{1}$$

where $I_{\mathrm{pred}}$ is the model's prediction for the measured intensity, $\mathbf{h}_i$ is the index of the symmetry-equivalent reflection in the asymmetric unit of reciprocal space, and $a$, $b$, $c$, and $d$ are functions of the experimental geometry; $\phi_i$ is the spindle rotation angle, $s_i = |\mathbf{s}_i|$ is the scattering vector magnitude, $p_i$ is the detector chip index, and $(x_i, y_i)$ is the position in the detector plane. Roughly speaking, $a$ corrects for absorption, $b$ corrects for overall changes in illuminated volume and beam intensity, $c$ is strictly positive and corrects for excess isotropic scattering (which may occur if extra material, such as the crystal loop, passes through the beam), and $d$ corrects for detector chip efficiency (flat-field errors). The continuous functions $a$, $b$, and $c$ were obtained by linear interpolation on multi-dimensional grids. A $9 \times 9$ grid was used for the detector plane position, 100 grid points were used for scattering vector ($0 < s < 0.9132\,\text{Å}^{-1}$), and 26 grid points were used for the spindle angle coordinate of each 50° data wedge. A set of 960 discrete values was used for $d$, corresponding to the 960 detector chips in the Pilatus 6M.

The parameters of the scaling model were fit by minimizing the sum of the $\chi^2$ and regularization terms, as follows:

$$\mathcal{H} = \sum_i \left( I_{\mathrm{meas}}(i) - I_{\mathrm{pred}}(i) \right)^2 \sigma_i^{-2} + \sum_j \lambda_j \mathcal{B}_j, \tag{2}$$

where $\sigma_i$ is the uncertainty (standard error) estimate for $I_{\mathrm{meas}}(i)$, $\mathcal{B}_j$ are the regularization functions and $\lambda_j$ are the corresponding weights (Lagrange multipliers). The regularization functions are used to stabilize refinement and to enforce smoothness of the correction factors. For the correction factors $a$, $b$, and $c$, smoothness was enforced by minimizing the second derivative. Discrete approximations (*11*) of the following integrals were used: $\int d\phi dx dy \left| \partial_\phi^2 a \right|^2$,

$\int d\phi dxdy \left|\partial_x^2 a + \partial_y^2 a\right|^2$, $\int d\phi \left|\partial_\phi^2 b\right|^2$, $\int d\phi ds \left|\partial_\phi^2 c\right|^2$, $\int d\phi ds \left|\partial_s^2 c\right|^2$. In addition, the offset correction was forced to be positive, and to stabilize the refinement, its magnitude was minimized using a discrete approximation of $\int d\phi ds \left|c\right|^2$. Finally, the detector correction factors were regularized using $\sum_p |d(p) - 1|^2$, which ensures $d = 1$ in the absence of data. The nonlinear minimization problem was solved iteratively by alternately minimizing $\mathcal{H}$ with constant $I_{\text{merge}}$ (a linear problem) and updating $I_{\text{merge}}$ given the new scale factors (*12*). To simplify the implementation, each set of parameters was refined individually (or in pairs) with the others held fixed. Satisfactory results were obtained by refining corrections in the following sequence: $\{b, o, cb, o, c, a, d\}$, where $cb$ refers to fitting the model for $c$ followed by $b$, and $o$ is an outlier rejection step (Figures S6 and S7).

After refining the scaling model, redundant observations in the continuous scattering map were merged. Observations more than $5\sigma$ from the mean were excluded. The estimated Bragg intensities, obtained from integration of the masked voxels, were also scaled and merged using the same model, except that the offset correction was omitted and an outlier cutoff of $2\sigma$ was used. The merged values were compared with the Bragg intensities integrated and merged using xds (*4*) and aimless (*5*). A single scale factor was found to bring the xds/aimless values into agreement with our Bragg intensity map. Since the intensities determined by xds and aimless are more accurate and complete than our estimates, the xds/aimless values were used instead for all subsequent analysis. Doing so also ensures that the Bragg intensities matched the values used for structure determination.

**Placement of intensities on an absolute scale.** After merging, the intensities were placed on an absolute scale. The overall scale factor was found by adaptation of the total intensity method originally described by Krogh-Moe (K-M) (*13, 14*). The standard K-M method, described in Appendix 2 in the Supplementary Text, involves predicting the contribution of individual atoms to the total intensity ($I_{\text{total, predicted}}$, Equation 60) and comparing the prediction to the measured value ( $I_{\text{total, measured}}$, Equation 61) to determine a scale factor $\alpha$ as follows:

$$\alpha = \frac{I_{\text{total, predicted}}}{I_{\text{total, measured}}}. \tag{3}$$

To test for convergence, the scaling factor was calculated in two ways: first using the standard K-M method, and second using a modified K-M method to account for inter-atomic interference (Figure S8). For the standard K-M scaling method, an estimate of the atomic inventory of the unit cell was used to calculate the theoretical coherent and incoherent scattering for independent atoms. The theoretical scattering calculation included 290 water molecules, 6 nitrate ions, and 1 lysozyme molecule in the unit cell, for a total of 1546 H, 613 C, 199 N, 493 O, and 10 S atoms. The total number of electrons was $Z = 10720$. Then, $I_{\text{total, predicted}}$ was calculated using Equation 60, integrating over the observed region of reciprocal space. The modified K-M method was identical to the standard K-M method, except that $I_{\text{total, predicted}}$ was modified to include the interference between all atom pairs whose average inter-atomic distance could be predicted from the chemical structure alone (i.e. the protein sequence and known structure of water and solutes). Both covalent bonding and torsional restraints were considered. Molecular coordinates were taken from the chemical component dictionary (*15*), and pair distances between atoms of adjacent amino acids in the sequence were calculated assuming a planar peptide bond with a bond length of 1.33Å. This resulted in 7405 pair distances for lysozyme, 6 per nitrate molecule, and 3 per water molecule. Then, the following bonding correction was calculated and added to the elastic scattering in Equation 60:

$$I_{\text{bond}}(s) = 2\sum_{n,m} f_n(s) f_m(s) \frac{\sin(2\pi s\, r_{nm})}{2\pi s\, r_{nm}}, \tag{4}$$

where $f$ is the atomic scattering factor, the sum is over bonded atom pairs, and $r_{nm}$ is the inter-atomic distance.

With synthetic data, both methods converge to the expected value of 1 for $\alpha$ (Fig. S8, left). However, the modified method converges much more quickly and provides a more accurate scale factor at the resolutions ($\sim 2\,\text{Å}$) that are typical for macromolecular crystallography.

**Generation of the final diffuse map.** A final map of the diffuse intensities $I_D$ was generated on a fine grid with 13, 11, and 11 subdivisions along the reciprocal unit cell vectors $\mathbf{a}^*$, $\mathbf{b}^*$, and $\mathbf{c}^*$, respectively. The resolution range of the map was $25 - 1.25\,\text{Å}$ (scattering vector of $0.04 - 0.8\,\text{Å}^{-1}$). Voxels containing Bragg peaks were excluded. Geometric and background corrections were applied (Equation 52), and redundant observations were merged using the scaling model derived from the coarse map, described above. Errors were estimated using Poisson statistics and propagated through the correction, scaling and merging steps. When merging, observations with intensities more than $5\sigma$ from the mean were flagged as outliers and excluded. Intensities were placed on an absolute scale using the previously-determined scale factor $\alpha$, and the theoretical incoherent scattering was subtracted (Equation 56). To calculate the correlation coefficient for random half-datasets, the unmerged observations were randomly assigned using an algorithm that gave approximately equal statistical weight to each half-dataset, and the half-datasets were merged separately.

#### All-atom molecular dynamics (MD) simulation

Four all-atom MD simulations of triclinic lysozyme crystals were performed with 1, 27 (3x3x3), 125 (5x5x5), and 343 (7x7x7) unit cells, similar to a 12 unit-cell simulation described previously (*16*). The simulation was prepared using the AMBER suite, version 18 (*17*), using the ff14SB force field for the protein (*18, 19*), the SPC/E model for water (*20*), and the general Amber force field (GAFF) (*21*) parameters for the nitrate ion. The simulation boxes had dimensions equal to integer multiples of the experimentally determined room-temperature unit cell from PDB ID *4lzt* ($a = 27.24\,\text{Å}$, $b = 31.87\,\text{Å}$, $c = 34.23\,\text{Å}$, $\alpha = 88.52°$, $\beta = 108.53°$, $\gamma = 111.89°$). The simulation was initialized with the measured protein atom coordinates from PDB ID *4lzt* (using the "A" alternate conformer) (*8*), arranged in a supercell grid. Nine nitrate ions were added per unit cell to neutralize the charge, as well as 290 water molecules. The number of water molecules was manually adjusted in order to achieve $\sim 1$ atm pressure at 295 K, resulting in 290, 293, 284, and 270 waters per protein chain in the 1, 27, 125, and 343 unit cell simulations, respectively. The simulations were equilibrated for about 0.2 $\mu$s and continued for an additional 5, 5, 2 and 1 $\mu$s, respectively, saving coordinates every 0.4 ns. A time step of 4 fs was used, where non-water hydrogen masses are set to 3 amu, with a corresponding decrease in the mass of its bonded atom (*22*).

For each snapshot, structure factors were calculated from the atomic coordinates using the ccp4 (*6*) program *sfall* with its default grid parameters, a resolution of 0.95 Å, and a VDWR parameter of 3.0. The B-factor was set to $15\,\text{Å}^2$ for each atom. This value for a "snapshot" B-factor smooths the electron density distribution to allow the Fourier transforms used by *sfall* to obtain a converged result; this was tested by comparing to test calculations using twice as many grid points in each dimension and for test calculations in which the "snapshot" B-factor was varied between 5 and $20\,\text{Å}^2$. Since every atom was assigned the same B-factor, its effect can be undone by multiplying the structure factors coming from the *sfall* run by $\exp\left(+Bs^2/4\right)$. The Bragg intensity per unit cell (Equation 36) then was calculated using $I_B = N^{-2}\left\langle F(h_0)\right\rangle^2$ where $F(h_0)$ is the supercell structure factor evaluated at the Bragg positions $h_0$, $N$ is the number of unit cells, and brackets represent an average over all saved simulation frames (see Appendix 1 in the Supplementary Text). Similarly, the diffuse scattering per unit cell (Equation 35) was calculated using $I_D = N^{-1}(\left\langle F^2\right\rangle - \left\langle F\right\rangle^2)$. The whole procedure is encapsulated in the *md2diffuse.sh* script, distributed as a part of the AmberTools distribution (`http://ambermd.org`).

#### Lattice dynamics simulation

Lattice dynamics simulations and model refinement were performed in Matlab. Protein molecules were modeled as rigid bodies, and the lattice contacts were modeled as an elastic network (*23–27*) with pair-wise interactions between $\alpha$ carbons. The lattice contacts were identified in the all-atom structure determined in this study (PDB ID *6o2h*). First, atoms with alternate conformers were assigned to their occupancy-weighted average positions. Then, the atomic coordinates of the 26 nearest neighbors in the lattice were generated by applying the crystal symmetry operators. A lattice contact was defined between any atom in the central protein chain that came within 4 Å of an atom belonging to a neighbor. Finally, the network was reduced to a $C_\alpha$ model. If any atoms belonging to a pair of residues formed a lattice contact, a spring was created between the $C_\alpha$ atoms in the network. A total of 100 intermolecular springs were modeled, of which 50 were unique due to crystal symmetry.

Two types of pair potential were modeled: *Gaussian* and *directional*, as follows:

$$V_{jj'}^{(\text{Gauss.})} = \frac{1}{2}\gamma_{jj'}\left|\mathbf{u}_{(j)} - \mathbf{u}_{(j')}\right|^2 \tag{5}$$

and

$$V_{jj'}^{(\text{dir.})} = \frac{1}{2}\gamma_{jj'}\left(\left(\mathbf{u}_{(j)} - \mathbf{u}_{(j')}\right)\cdot\hat{\mathbf{r}}_{(j,j')}\right)^2, \tag{6}$$

where $j$ and $j'$ are the node indices, $\mathbf{u}$ is the displacement vector of a node from its equilibrium position, $\gamma$ is a spring constant, and $\hat{\mathbf{r}}_{(j,j')}$ is the unit vector pointing from node $j$ to $j'$.

The equations of motion were solved in a rigid-body vibrational coordinate system using the Born/Von-Karman method (Appendix 3 in the Supplementary Text). The diffuse scattering was calculated for a 13x11x11 periodic supercell, chosen to match the level of detail in the experimental map, using the one-phonon approximation (Equation

82). Terms in the one-phonon structure factor (Equation 83) were calculated using the fast Fourier transform-based method (*28*) with form factors approximated by four Gaussians and a constant (*29, 30*). The scattering contribution from the mean solvent density was modeled using Babinet's principle: since any constant can be added to the electron density without changing the structure factor (except at $\mathbf{s} = 0$), a constant density of $\rho_{\text{solv.}}$ surrounding a protein can be equivalently modeled by a density of 0 and $-\rho_{\text{solv.}}$ in the solvent-excluded region. For reasons of computational convenience, the excluded solvent can then be represented by pseudo-atoms with Gaussian form factors. To calculate the Babinet representation, voxels of the excluded solvent mask from REFMAC5 (*9*) were divided among the modeled atoms based on proximity. The set of voxels associated with each atom was approximated by a three-dimensional anisotropic Gaussian with the same first and second moments of density. The overall solvent scaling parameters $k_{\text{solv.}}$ and $B_{\text{solv.}}$ were then adjusted to minimize the least-squares difference between $F_{\text{obs.}}$ and $|F_{\text{model}}|$, defined as follows:

$$F_{\text{model}} = F_{\text{calc.}} + k_{\text{solv.}} \exp(-B_{\text{solv.}} s^2/4)\, F_{\text{solv.}}, \tag{7}$$

where $F_{\text{calc.}}$ is the structure factor of the modeled atoms (protein and ordered solvent) and $F_{\text{solv.}}$ is the structure factor of the excluded solvent. The refined parameters $k_{\text{solv.}}$ and $B_{\text{solv.}}$ were then applied to the excluded-solvent form factors. The resulting pseudo-atoms were included in the list of atoms occupying the unit cell and assigned to the same rigid group as the nearest protein atom.

Spring constants in the model were refined in order to minimize the least-squares difference between the simulated one-phonon scattering and the measured variational scattering around the 400 most intense halos in the 2-2.5 Å resolution range. The reduced $\chi^2$ was calculated as follows:

$$\chi^2_{\text{red.}} = \left(\sum_{n=1}^{N} M_n\right)^{-1} \sum_{n=1}^{N} \sum_{m=1}^{M_n} \left(\frac{I^{(\text{meas.})}_{n,m} - I^{(\text{calc.})}_{n,m} - b_n}{\sigma_{n,m}}\right)^2, \tag{8}$$

where $N = 400$ is the number of halos fit, $M_n$ is the number of measured voxels around the $n^{\text{th}}$ halo (typically $M_n = 13 \times 11 \times 11 - 1 = 1572$), $b_n$ is an arbitrary constant offset for each halo (determined separately by least-squares minimization for each $n$), and $\sigma$ is the experimental uncertainty. The spring constants were refined in four stages. In the first stage, all springs were set to Gaussian springs and assigned the same spring constant. In the second stage, springs belonging to the same interface (those involving a particular neighbor) were given the same spring constant. In the third stage, the pair-potentials for each interface were allowed to be a linear combination of Gaussian and directional. Finally, each pair potential was refined individually with a linear combination of Gaussian and directional springs. The overall $\chi^2$ was monitored during refinement to assess whether adding the extra degrees of freedom to the model significantly improved the fit (Fig. S13).

After refining the model, the scattering was calculated throughout reciprocal space using the one-phonon approximation (Equation 82). The Pearson correlation coefficient (CC) between the measured variational scattering map and the simulation was calculated within shells of constant resolution spanning $0.04\,\text{Å}^{-1}$ to $0.80\,\text{Å}^{-1}$ with a constant width of $\Delta s = 0.02\,\text{Å}^{-1}$ . Within each resolution bin, CC was calculated as follows:

$$\text{CC} = \frac{\sum_n \left(I_{\text{meas.}}(\text{s}_n) - \bar{I}_{\text{meas.}}\right)\left(I_{\text{calc.}}(\text{s}_n) - \bar{I}_{\text{calc.}}\right)}{\sqrt{\sum_n \left(I_{\text{meas.}}(\text{s}_n) - \bar{I}_{\text{meas.}}\right)^2 \sum_n \left(I_{\text{calc.}}(\text{s}_n) - \bar{I}_{\text{calc.}}\right)^2}}, \tag{9}$$

where $\bar{I}_{\text{meas.}}$ and $\bar{I}_{\text{calc.}}$ are the mean intensities in that resolution bin, and the sums are over all measured voxels within the resolution bin (Fig. S14). For comparison with the MD simulation, which was calculated on a coarser 7x7x7 sub-sampled reciprocal lattice, the full map was interpolated at the voxels of the coarser grid by least-squares fitting a 2nd order polynomial over all neighboring voxels (a 3x3x3 kernel). Voxels at the Bragg positions (those with integer Miller indices) were excluded. The CC was calculated, as above, between the interpolated simulated map and a similarly interpolated experimental map (Fig. 2D in the Main Text).

#### Internal protein dynamics simulation

Internal dynamics simulations and model refinement were performed in Matlab. The dynamics of lysozyme within the crystal environment were simulated using an all-atom elastic network where each residue was restrained to move as a rigid body, and lattice contacts were explicitly modeled. To generate the model, first the atoms with alternate conformers were assigned to their occupancy-weighted average positions. Then, springs were created between any pair of non-H protein atoms belonging to different residues within a cutoff distance of 4 Å. Intermolecular springs were modeled between atoms in the protein and those of its neighbors in the lattice within the 4 Å cutoff distance. All springs were of the directional type (Equation 6).

The equations of motion for a single unit cell were solved using the Born/Von-Karman method as described for the lattice dynamics simulation (Appendix 3 in the Supplementary Text), except that the potential energy function was modified in order to remove those modes associated with rigid-body motion of the entire protein. This was done by assigning the component of displacement associated with such motions a restoring force of zero. The normal modes associated with rigid-body displacements then have eigenvalues of zero and are eliminated during generalized inversion of the dynamical matrix (discussed in Appendix 3 in the Supplementary Text). More specifically, components of the Hessian matrix (Equation 65) were modified as follows:

$$\mathbf{V}_{(l,l')} \coloneqq \mathbf{P}^{\mathbf{T}}\mathbf{V}_{(l,l')}\mathbf{P}, \tag{10}$$

where $\mathbf{P}$ is an operator that projects out the rigid-body component of displacement,

$$\mathbf{P} = \mathbf{I} - (\mathbf{A}\backslash\mathbf{A}_0)(\mathbf{A}_0\backslash\mathbf{A}), \tag{11}$$

$\mathbf{I}$ is a $6m \times 6m$ identity matrix ($m = 129$ is the number of residues), $\mathbf{A}$ is a $3n \times 6m$ matrix ($n$ is the number of atoms in the protein) that transforms between the Cartesian atomic displacement coordinates, $\mathbf{u}$, and the generalized coordinates of the internal dynamics model (Equation 63), $\mathbf{A}_0$ is a $3n \times 6$ matrix that transforms between $\mathbf{u}$ and the generalized coordinates of the lattice dynamics model, and the forward slash signifies left matrix division (if $\mathbf{X} = \mathbf{A}\backslash\mathbf{A}_0$, then $\mathbf{X}$ is the least squares solution to the system of equations $\mathbf{AX} = \mathbf{A}_0$).

The model was parameterized with one coupling constant per residue, so that a spring connecting a pair of atoms ($j$ and $j'$) has a spring constant equal to the geometric mean of the residues' coupling constants $g_i$ and $g_{i'}$, as follows:

$$\gamma_{j,j'} = \sqrt{g_i g_{i'}}. \tag{12}$$

The parameters were optimized in order to minimize the $\chi^2$ between the measured and simulated atomic displacement parameters (ADPs), calculated as follows:

$$\chi^2 = \sum_{j=1}^{N}\sum_{n=1}^{9}\left(\left(\mathbf{U}_j^{(\text{meas.})}\right)_n - \left(\mathbf{U}_j^{(\text{latt.})} + \mathbf{U}_j^{(\text{int.})}\right)_n\right)^2, \tag{13}$$

where $(\mathbf{U}_j)_n$ is the $n^{\text{th}}$ component of the ADP for atom $j$ ($\mathbf{U}_j$ is a symmetric 3x3 matrix with 9 components), $\mathbf{U}_j^{(\text{latt.})}$ is the calculated ADP for the fully-refined lattice dynamics model, and $\mathbf{U}_j^{(\text{int.})}$ is the calculated ADP for the internal dynamics model (Equation 76).

After refining the model, the displacement correlations were assessed using the directional correlation coefficient, defined as follows:

$$\text{CC}_{j,j'} = \frac{\hat{\mathbf{r}}_{j,j'}^{\text{T}}\left\langle \mathbf{u}_j \mathbf{u}_{j'}^{\text{T}}\right\rangle \hat{\mathbf{r}}_{j,j'}}{\sqrt{(\text{Tr}\mathbf{U}_j/3)(\text{Tr}\mathbf{U}_{j'}/3)}}, \tag{14}$$

where $\hat{\mathbf{r}}_{j,j'}$ is the unit vector pointing from atom $j$ to $j'$.

We also defined an alternate model of internal protein motion where the modes associated with rigid-body displacements of individual domains are suppressed. Residues were assigned to three domains (*31*) as follows: 5-36 and 98-129 to $\alpha$, 40-94 to $\beta$, and those remaining to the hinge region. The Hessian matrix was modified as described above, except that the $\mathbf{P}$ operator appearing in Equation 10 was calculated as follows:

$$\mathbf{P} = \mathbf{I} - (\mathbf{A}\backslash\mathbf{A}_1)(\mathbf{A}_1\backslash\mathbf{A}), \tag{15}$$

were $\mathbf{A}_1$ is a $3n \times 6d$ matrix ($d = 3$ is the number of domains) that projects from the generalized coordinate system of the 3-domain model to the atomic displacements (Equation 63). The model was parameterized and refined as described above for the unrestrained model, and the directional correlation was calculated using Equation 14.

#### Diffuse Patterson map calculation

Diffuse Patterson maps were calculated in Matlab as the Fourier transform of the diffuse scattering (see Equation *41* and *Appendix 1* in the Supplementary Text), using a three-dimensional fast-Fourier transform (FFT).

The experimental diffuse map was pre-processed before performing the FFT to compensate for missing data. First, missing voxels in the diffuse map were filled in with the mean values from neighboring voxels. Then, the mean intensity in each resolution shell was subtracted, and voxels beyond the resolution limit of the map were filled with zeros. Finally, the data array was zero-padded to yield a diffuse Patterson map with a real-space voxel approximately 0.3 Å on a side (the voxel dimensions were $a/91$, $b/107$, and $c/115$, where $a$, $b$, and $c$ are the lattice constants).

The diffuse Patterson map for the refined vibrational model (lattice + internal) was calculated without approximation in the central region where $r < 25$ Å. To perform the calculation efficiently, the scattering per unit cell (79 in the Supplementary Text) was rearranged to single out a reference unit cell ($l = 0$):

$$I_D = \sum_j f_j \left\{ \sum_{l',j'} f_{j'} e^{2\pi i \mathbf{s}\cdot(\mathbf{r}_j - \mathbf{r}_{j'} - \mathbf{r}_{l'} + \mathbf{r}_0)} T_j T_{j'} \left( T_{0j,l'j'} - 1 \right) \right\}, \tag{16}$$

where the first sum runs over all atoms in the unit cell, $f_i$ is the atomic scattering factor, $T_j$ is the Debye-Waller factor (Equation 80), and $T_{0j,l'j'}$ depends on the cross-terms of the covariance matrix (Equation 81 with $l = 0$). The term in the curly brackets resembles the standard structure factor equation for the primed atoms, except that the origin is shifted and the Debye-Waller factor is replaced by

$$(T_{\text{eff}})_{0j,l'j'} = T_j T_{j'} \left( T_{0j,l'j'} - 1 \right). \tag{17}$$

The effective Debye-Waller factor was separated into contributions from lattice and internal motion:

$$T_{\text{eff}} = T_{\text{eff}}^{\text{latt}} + T_{\text{eff}}^{\text{int}}. \tag{18}$$

The lattice term was calculated as follows:

$$\left( T_{\text{eff}}^{\text{latt}} \right)_{0j,l'j'} = T_j T_{j'} \left( T_{0j,l'j'}^{\text{latt}} - 1 \right), \tag{19}$$

where the experimentally determined ADPs were used in $T_j$ and $T_{j'}$, and the remaining $T$ was calculated from the refined lattice model (Equations 75 and 80). This corresponds to the definition used in the one-phonon simulation (Equation 82). For the internal motions, the effective Debye-Waller factor was calculated as follows:

$$\left( T_{\text{eff}}^{\text{int}} \right)_{0j,l'j'} = T_j^{\text{latt}} T_j^{\text{int}} T_{j'}^{\text{latt}} T_{j'}^{\text{int}} T_{0j,l'j'}^{\text{latt}} \left( T_{0j,l'j'}^{\text{int}} - 1 \right), \tag{20}$$

where the $T$'s are calculated from the covariance matrices of the lattice and internal dynamics simulations.

Excluded solvent effects were modeled by pseudo-atoms with Gaussian scattering factors, as described above for the lattice dynamics simulation. In Equation 16, terms in curly brackets were calculated using the FFT-based method as described for the lattice dynamics simulation, except that each atom had an effective Debye-Waller factor (Equation 18) and coordinates relative to $\mathbf{r}_j$. Since only the central part of the Patterson was desired, the sum was carried out over all atoms in the unit cell and its 26 nearest neighbors that satisfied $|\mathbf{r}_j - \mathbf{r}_{j'} - \mathbf{r}_{l'} + \mathbf{r}_0| < 29$ Å (the cutoff distance was chosen to be somewhat larger than the maximum distance of 25 Å to avoid truncation artifacts). After calculating the diffuse intensity map using Equation 16, the mean intensity in each resolution shell was subtracted and voxels outside the experimental resolution limit of 1.25 Å were set to zero. Then, the map was zero-padded, and the Patterson function was calculated using the three-dimensional FFT, as described above for the experimental map.

The reciprocal space correlation coefficients between diffuse Patterson maps were also calculated in Matlab. First, real space voxels with $|\mathbf{r}| < 2$ Å or $|\mathbf{r}| > 25$ Å were set to zero, and maps were truncated at $|x| < a$, $|y| < b$ and $|z| < c$ so that the reciprocal space map would be oversampled by a factor of 2 in each direction. Then, the inverse FFT of each truncated map was calculated. The Pearson correlation coefficients (Equation 9) between the experimental and simulated intensity maps were calculated in shells of constant resolution spanning 0.04 to 0.80 $\text{Å}^{-1}$ with bin widths of $\Delta s = 0.04\,\text{Å}^{-1}$ .

### Supplementary Text

#### Appendix 1 General Theory

The kinematical theory of X-ray diffraction applicable to diffuse scattering from proteins has been described previously (*32, 33*). Here we provide an overview of the main results as relevant to this study.

**Total scattering cross-section.** The macroscopic differential scattering cross-section, $d\Sigma/d\Omega$, describes the probability of observing a scattering event in the solid angle $d\Omega$ when the sample is illuminated by X-rays. The cross-section depends on the respective wavevectors of the incident and scattered X-rays, $\mathbf{s}_0$ and $\mathbf{s}'$. The wavevectors have magnitudes of inverse wavelength: $s_0 = |\mathbf{s}_0| = 1/\lambda$ and $s' = |\mathbf{s}'| = 1/\lambda'$, where $\lambda$ and $\lambda'$ are the wavelengths of the incident and scattered X-ray photons. Given an incident flux, $J_0$, the total scattered flux into a solid angle $\Delta\Omega$ in the direction $\hat{s}'$ is as follows:

$$J(\hat{\mathbf{s}}') = J_0 \Delta\Omega \frac{d\Sigma}{d\Omega}(\mathbf{s}_0, \mathbf{s}'). \tag{21}$$

In the X-ray experiments described here, both coherent and incoherent processes contribute to X-ray scattering:

$$\frac{d\Sigma}{d\Omega} = \left(\frac{d\Sigma}{d\Omega}\right)_{\text{coherent}} + \left(\frac{d\Sigma}{d\Omega}\right)_{\text{incoherent}}. \tag{22}$$

The coherent X-ray scattering has a cross-section

$$\left(\frac{d\Sigma}{d\Omega}\right)_{\text{coherent}} = I_e(\hat{\mathbf{s}}_0, \hat{\mathbf{s}}') I_{\text{sample}}(\mathbf{s}), \tag{23}$$

where $\mathbf{s} = \mathbf{R}^{\text{T}}(\mathbf{s}' - \mathbf{s}_0)$ is the scattering vector in the sample frame which is rotated with respect to the lab frame by a rotation matrix $\mathbf{R}$, $I_{\text{sample}}$ is the coherent intensity of the sample in per-electron units, and $I_e$ is the Thomson scattering cross-section of a free electron,

$$I_e = r_e^2 P(\hat{\mathbf{s}}_0, \hat{\mathbf{s}}'), \tag{24}$$

where $r_e$ is the classical electron radius and $P(\hat{\mathbf{s}}_0, \hat{\mathbf{s}}')$ is the polarization factor.

Incoherent, or Compton, X-ray scattering has a cross-section (*34*):

$$\left(\frac{d\Sigma}{d\Omega}\right)_{\text{incoherent}} = I_e(\hat{\mathbf{s}}_0, \hat{\mathbf{s}}') S_{\text{sample}}(\mathbf{s}), \tag{25}$$

where $S_{\text{sample}}$ is the sum of atomic incoherent scattering functions over all illuminated atoms (*29*). The Compton scattering process is inelastic: the scattered photon has less energy than the incident photon by an amount that increases with scattering angle. However, the wavelength shift, which is at most twice the Compton wavelength of the electron $\lambda_{\text{C}} = h m_e^{-1} c^{-1} \approx 0.02426\,\text{Å}$, is small compared with the $\lambda \sim 1\,\text{Å}$ wavelengths generally employed in X-ray crystallography. Therefore, in the following, we assume $\lambda = \lambda'$.

The sample in this case is a crystal composed of unit cells. The crystal is assumed to be homogeneous, meaning that the unit cells are equivalent to each other on average, and therefore one can write:

$$I_{\text{sample}}(\mathbf{s}) = (V_0/v_c) I(\mathbf{s}) \tag{26}$$

and

$$S_{\text{sample}}(\mathbf{s}) = (V_0/v_c) S(\mathbf{s}), \tag{27}$$

where $V_0$ is the volume of the sample illuminated by the X-ray beam, $v_c$ is the unit cell volume ($V_0/v_c$ is the number of unit cells illuminated), and $I(\mathbf{s})$ and $S(\mathbf{s})$ are the coherent and incoherent intensities *per unit cell.* Finally, we can write the scattered flux in terms of the intensities per unit cell by combining Equations 21-27, as follows:

$$J(\hat{\mathbf{s}}') = J_0 \Delta\Omega r_e^2 P(\hat{\mathbf{s}}_0, \hat{\mathbf{s}}')(V_0/v_c)\left[I(\mathbf{s}) + S(\mathbf{s})\right]. \tag{28}$$

**Coherent scattering from crystals.** Coherent scattering of X-rays depends on the spatial distribution of electrons in the sample. If $\rho(\mathbf{r})$ is the instantaneous number density of electrons at a position $\mathbf{r}$, the scattering is described by the Fourier transform of the electron density, a quantity known as the structure factor, $F(\mathbf{s})$:

$$F(\mathbf{s}) = \int \rho(\mathbf{r}) e^{2\pi i \mathbf{s}\cdot\mathbf{r}} d^3\mathbf{r}. \tag{29}$$

The intensity per unit cell is proportional to the mean square structure factor, as follows:

$$I(\mathbf{s}) = N^{-1} \left\langle F(\mathbf{s})^2 \right\rangle, \tag{30}$$

where $N$ is the number of unit cells under consideration and brackets denote the ensemble (or time) average. In general, the electron density is time-dependent, but the system is in equilibrium so that we can define an average electron density $\langle \rho(\mathbf{r}) \rangle$ with corresponding structure factor $\langle F(\mathbf{s}) \rangle$. If the structure factor is separated into the average and fluctuating terms, $F = \langle F \rangle + \tilde{F}$, the total intensity (Equation 30) becomes

$$I(\mathbf{s}) = N^{-1} \langle F(\mathbf{s}) \rangle^2 + N^{-1} \left\langle \tilde{F}(\mathbf{s})^2 \right\rangle. \tag{31}$$

We can further define an average unit-cell electron density, in which the densities of all unit cells are superimposed, $\langle \rho_{\text{cell}}(\mathbf{r}) \rangle$, and corresponding structure factor $\langle F_{\text{cell}}(\mathbf{s}) \rangle$. If the unit cells under consideration are equivalent, ensemble averaging is equivalent to spatial averaging, and the two average structure factors are related as follows:

$$\langle F(\mathbf{s}) \rangle = \langle F_{\text{cell}}(\mathbf{s}) \rangle \sum_{\mathbf{n}} e^{2\pi i \mathbf{s}\cdot\mathbf{r_n}}, \tag{32}$$

where the sum runs over all unit cells, $N$, and $\mathbf{r_n}$ is the origin of each unit cell. If $\mathbf{s}$ happens to coincide with a node of the reciprocal lattice $\mathbf{g_h}$ ($\mathbf{h}$ is the vector of Miller indices, $h$, $k$, and $l$), the phase factor in the sum is equal to 1 for all $\mathbf{n}$, and the sum evaluates to $N$. When $N$ is large, the amplitude of the lattice term falls off very quickly with $|\mathbf{s} - \mathbf{g_h}|$ , and as $N \to \infty$, it can be written as a sum of delta functions as follows:

$$\lim_{N\to\infty} \left| \sum_{\mathbf{n}} e^{2\pi i \mathbf{s}\cdot\mathbf{r_n}} \right|^2 = N v_c^* \sum_{\mathbf{h}} \delta(\mathbf{s} - \mathbf{g_h}), \tag{33}$$

where $v_c^*$ is the volume of a unit cell of the reciprocal lattice. Thus, for the large crystal, we can combine Equations 31-33 to write the intensity per unit cell as follows:

$$I(\mathbf{s}) = \langle F_{\text{cell}}(\mathbf{s}) \rangle^2 \sum_{h} v_c^* \delta(\mathbf{s} - \mathbf{g_h}) + N^{-1} \left\langle \tilde{F}(\mathbf{s})^2 \right\rangle. \tag{34}$$

The second term of Equation 34 is the diffuse scattering per unit cell,

$$\begin{aligned} I_D(\mathbf{s}) &= N^{-1} \left\langle \tilde{F}(\mathbf{s})^2 \right\rangle \\ &= N^{-1} \left( \left\langle F(\mathbf{s})^2 \right\rangle - \langle F(\mathbf{s}) \rangle^2 \right). \end{aligned} \tag{35}$$

The first term of Equation 34 contains the Bragg peaks. We can define the Bragg intensity as follows:

$$I_B(\mathbf{s}) = \langle F_{\text{cell}}(\mathbf{s}) \rangle^2, \tag{36}$$

which is obtained for $\mathbf{s} = \mathbf{g_h}$ upon integration of the Bragg peaks in reciprocal space. With these definitions (Equations 35-36), the coherent intensity from a macroscopic crystal (Equation 34) is as follows:

$$I(\mathbf{s}) = I_D(\mathbf{s}) + I_B(\mathbf{s}) \sum_{h} v_c^* \delta(\mathbf{s} - \mathbf{g_h}). \tag{37}$$

**Diffuse Patterson function.** The diffuse Patterson function, or three-dimensional difference pair distance distribution function (3D-$\Delta$PDF), is defined as the mean autocorrelation of the electron density fluctuations per unit cell, as follows:

$$\begin{aligned} \Delta\text{PDF}(\mathbf{r}) &= N^{-1} \langle \Delta\rho \star \Delta\rho \rangle (\mathbf{r}) \\ &= N^{-1} \int \langle \Delta\rho(\mathbf{u}) \Delta\rho(\mathbf{r} + \mathbf{u}) \rangle \, d^3\mathbf{u}, \end{aligned} \tag{38}$$

where $\Delta\rho = \rho - \langle\rho\rangle$ is the difference between the instantaneous electron density and the ensemble average. The diffuse intensity per unit cell (Equation 35) can be rearranged as:

$$I_D(\mathbf{s}) = N^{-1}\left\langle |F(\mathbf{s}) - \langle F(\mathbf{s})\rangle|^2 \right\rangle . \tag{39}$$

Plugging in the definition of the structure factor (Equation 29), we see that $I_D$ is the Fourier transform of the $\Delta$PDF, as follows:

$$\begin{aligned} I_D(\mathbf{s}) &= N^{-1}\left\langle \left| \int \Delta\rho(\mathbf{r}) e^{2\pi i \mathbf{s}\cdot\mathbf{r}} d^3\mathbf{r} \right|^2 \right\rangle \\ &= N^{-1} \int \langle \Delta\rho \star \Delta\rho\rangle (\mathbf{r})\, e^{2\pi i \mathbf{s}\cdot\mathbf{r}} d^3\mathbf{r} \\ &= \int \Delta\mathrm{PDF}(\mathbf{r})\, e^{2\pi i \mathbf{s}\cdot\mathbf{r}} d^3\mathbf{r}. \end{aligned} \tag{40}$$

Thus, $I_D$ and $\Delta$PDF are a Fourier transform pair, and the inverse transform is:

$$\Delta\mathrm{PDF}(\mathbf{r}) = \int I_D(\mathbf{s})\, e^{-2\pi i \mathbf{s}\cdot\mathbf{r}} d^3\mathbf{s}\, . \tag{41}$$

**Scattering simulation.** In order to simulate total scattering without having to model the entire crystal, it is useful to simulate a small number of unit cells and to impose periodic boundary conditions by what is known as the supercell method (*16, 35, 36*). Given a supercell consisting of a three-dimensional array of $N_{\mathrm{scell}} = N_1 \times N_2 \times N_3$ unit cells, a "supercell reciprocal lattice" can be defined, which subdivides the regular reciprocal lattice by $N_1$ along $\mathbf{a}^*$, $N_2$ along $\mathbf{b}^*$, and $N_3$ along $\mathbf{c}^*$. The total intensity per unit cell (Equation 30) is calculated in the same manner as usual (Equation 31), except that the structure factor $F_{\mathrm{scell}}$ is now a periodic function whose Fourier transform is defined only at nodes of the reciprocal lattice of the supercell. The mean of $F_{\mathrm{scell}}$, evaluated at the Bragg peak locations (the nodes of the unit cell reciprocal lattice, $\mathbf{g_h}$) is proportional to the regular unit cell structure factor $F_{\mathrm{cell}}$:

$$|\langle F_{\mathrm{scell}}(\mathbf{s} = \mathbf{g_h})\rangle| = N_{\mathrm{scell}}\, |\langle F_{\mathrm{cell}}(\mathbf{s})\rangle|\, . \tag{42}$$

Thus, we can calculate the Bragg intensity per unit cell (Equation 36) from a supercell simulation as follows:

$$I_B(\mathbf{s}) = N_{\mathrm{scell}}^{-2}\, \langle F_{\mathrm{scell}}(\mathbf{s})\rangle^2\, , \tag{43}$$

where $F_{\mathrm{scell}}$ is evaluated only at nodes of the regular, unit-cell reciprocal lattice. The diffuse intensity per unit cell is defined in the same way as the macroscopic crystal case (Equation 35), except that it is no longer a continuous function of $\mathbf{s}$ and is evaluated only at the nodes of the supercell reciprocal lattice.

### Appendix 2 Experimental Corrections

Here we consider geometric and experimental corrections that relate the photon counts observed by a detector to the coherent intensity per unit cell, $I(\mathbf{s})$ (see Appendix 1 and Equation 30). The first section describes the factors depending on the scattering geometry. The second section describes corrections to the Bragg intensity for a crystal that rotates continuously during the exposure. The final section describes a method for placing the total intensity on an absolute scale and subtracting the incoherent contribution.

**Geometric factors.** We consider corrections applicable to the geometry normally used in macromolecular crystallography, which consists of a planar X-ray detector with square pixels and a sample that is rotated about a fixed spindle axis. Corrections are derived assuming that detection occurs in the far field from a small sample, so that the illuminated portion of the sample acts as a point-like source.

In Appendix 1, Equation 28 relates the scattered flux $J(\hat{\mathbf{s}}')$ in the direction $\hat{\mathbf{s}}'$ to the coherent and incoherent intensities. Two prefactors in that equation, the polarization $P$ and solid angle $\Delta\Omega$, depend on the scattering geometry. The solid angle of the pixel can be calculated as follows:

$$\Delta\Omega = (\Delta x \Delta y / d^2)\cos\omega, \tag{44}$$

where $d$ is the distance between the sample and detector pixel, $\Delta x$ and $\Delta y$ are the pixel width and height, and $\omega$ is the angle between the detector surface normal and the path of the incident photon. The polarization factor depends on the scattering geometry as well as X-ray source properties: the fraction of polarization, $p$, and the vector normal to the plane of polarization, $\hat{\mathbf{n}}$. In terms of these parameters, the polarization factor can be written as follows (*37*):

$$P = p\left(1 + \left(\hat{\mathbf{s}}' \cdot \hat{\mathbf{s}}_0\right)^2\right) + (1-2p)\left(1 - \left(\hat{\mathbf{s}}' \cdot \hat{\mathbf{n}}\right)^2\right). \tag{45}$$

In addition to solid angle and polarization, we must also correct for the efficiency of detection. In an ideal situation, the expected number of photons recorded by a detector pixel, $n$, would be equal to the scattered flux (Equation 28) times the exposure time: $n = J\Delta t$. However, two non-ideal effects may be important. First is the possibility that the detector pixel does not absorb all of the photons it intercepts. For a pixel of thickness $\delta$ and absorption coefficient of the sensor $\kappa$, the fraction of photons absorbed, $E$, is as follows:

$$E = 1 - \exp(-\kappa\, \delta / \cos\omega). \tag{46}$$

Second is the possibility that the material between the sample and detector absorbs or scatters some of the X-rays before they can be detected. For example, if a uniform material (such as air) with attenuation coefficient $\mu$ occupies the space between the sample and the detector (the path length is $d$), the probability that a photon is transmitted, $A$, is as follows:

$$A = \exp(-\mu\, d). \tag{47}$$

The sample rotates during the exposure, so that each pixel samples a range of scattering vectors in the reference frame of the sample. Let $\hat{\mathbf{m}}$ be the spindle axis, $\phi$ the spindle angle, $\Delta t$ the exposure time, and $\Delta\phi/\Delta t$ the rate of rotation. Then, for small $\Delta\phi$ and $\Delta\Omega$, the volume element swept by the pixel in reciprocal space is as follows (*33*):

$$\Delta v^* = \Delta\phi\Delta\Omega\lambda^{-3}\left|\hat{\mathbf{m}} \cdot \left(\hat{\mathbf{s}} \times \hat{\mathbf{s}}'\right)\right|. \tag{48}$$

The measured intensity thus depends on the average flux (Equation 28) for this volume,

$$\langle J\rangle_{\Delta v^*} = J_0 \Delta\Omega r_e^2 P\,(V_0/v_c)\,\langle I + S\rangle_{\Delta v^*}, \tag{49}$$

where the brackets signify averaging over the volume element $\Delta v^*$, as follows:

$$\langle I + S\rangle_{\Delta v^*} \equiv \frac{1}{|\Delta v^*|}\int_{\Delta v^*} d^3\mathbf{s}\,(I(\mathbf{s}) + S(\mathbf{s})). \tag{50}$$

Finally, there may be other photons detected during the exposure that come from background sources, such as air scatter. The background scattering rate for each pixel, $r_b$, will be sample and instrument-dependent, but it can be estimated experimentally, for example by performing a measurement with the sample removed.

With all corrections put together, the expected number of photons detected is related to the intensity per unit cell as follows:

$$n = \Delta t\left(r_b + \Delta\Omega EAPJ_0 r_e^2\,(V_0/v_c)\,\langle I + S\rangle_{\Delta v^*}\right). \tag{51}$$

**Corrections to experimental data.** Equation 51 relates the measured photon counts to the signal of interest: the coherent intensity per unit cell, $I$, which consists of the Bragg and diffuse components. Here we describe the data processing steps that are necessary to convert photon counts into estimates of $I_B$ and $I_D$ on an absolute scale.

In a standard macromolecular crystallography experiment, intensity is determined on an arbitrary scale (this is because the incident flux, the illuminated volume, or both, are typically unknown), and the background scattering is not measured. This is adequate for Bragg intensity determination because the intensity is integrated relative to a local background, and the overall scale factor is fit during model refinement. For diffuse scattering, it is necessary to measure and subtract the background. As a first step in data analysis, the background and geometric corrections are applied to the measured photon counts, so that one obtains an intensity on an arbitrary scale, which we call $I_{\text{meas}}$, as follows:

$$I_{\text{meas}} \equiv \frac{n_{\text{meas}}/\Delta t - r_b}{\Delta\Omega EAP}, \tag{52}$$

where the factors in the denominator are defined in Equations 44-47. This intensity is proportional to true intensity per unit cell, with the expected theoretical relationship (Equations 51and 52):

$$I_{\text{meas}} \propto \langle I + S\rangle_{\Delta v^*}. \tag{53}$$

For a given exposure, the prefactor $\left(J_0 r_e^2 V_0/v_c\right)$ is a constant that is the same for all pixels.

The next step is to separate $I_{\text{meas}}$ into Bragg and continuous scattering components. This is simplest in the case where the coherent intensity $I$ has a diffuse component that varies gradually on the scale of $\Delta v^*$ and Bragg peaks that are delta function-like on this scale (Equation 37). Then, $I_{\text{meas}}$ can be treated differently depending on whether the integration volume $\Delta v^*$ contains a Bragg peak. If it does not, the intensity can be factored out of the integral (Equation 50), and we have $\langle I+S\rangle_{\Delta v^*} \approx I_D + S$. If the volume contains a Bragg peak (at $\mathbf{s}=\mathbf{g_h}$), we have $\langle I+S\rangle_{\Delta v^*} \approx I_D + S + (v_c^*/\Delta v^*)I_B$. Thus, the Bragg intensity can be estimated from $I_{\text{meas}}$ by first subtracting the continuous scattering beneath the Bragg peak and then scaling by $\Delta v^*/v_c^*$ (the Lorentz correction), as follows:

$$I_B \propto (\Delta v^*/v_c^*)\,(I_{\text{meas}} - I_{\text{bkg}}), \tag{54}$$

where $I_{\text{bkg}}$ is an estimate for the continuous scattering contribution at $\mathbf{h}$.

**Incoherent scattering and absolute intensity.** A convenient method for determining the absolute scale factor based on the integral of the total intensity was first described by Krogh-Moe (*13*) and further elucidated by Norman (*14*). The total intensity includes the coherent scattering (Bragg and diffuse) as well as incoherent Compton scattering. The total integrated intensity (per unit cell) is defined as follows:

$$I_{\text{total}} = \int d^3\mathbf{s}\,\left(I(\mathbf{s}) + S(\mathbf{s})\right), \tag{55}$$

where $I$ and $S$ are the coherent and incoherent scattering per unit cell. Compton scattering is insensitive to molecular structure and can be calculated given the atomic inventory of the unit cell as follows:

$$S = \sum_{\text{n}} S_n(s), \tag{56}$$

where the sum runs over all atoms in the unit cell and $S_n(s)$ are the atomic incoherent scattering functions (*29,34*). Application of Parseval's theorem from harmonic analysis allows us to write the integral of the elastic scattering in terms of independent contributions from each atom, as follows:

$$\int d^3\mathbf{s}\, I(\mathbf{s}) = \int_0^\infty 4\pi s^2 ds \sum_n f_n^2(s), \tag{57}$$

where $f_n$ are the coherent atomic scattering factors (*30*). Thus, both the coherent and incoherent contributions to the total intensity can be calculated from the atomic scattering functions, as follows:

$$I_{\text{total}} = \int_0^\infty 4\pi s^2 ds \sum_n \left(f_n^2(s) + S_n(s)\right). \tag{58}$$

Alternatively, the total intensity can be found by directly integrating the measured three-dimensional reciprocal space map. Care must be taken however to place Bragg and diffuse intensities on the same (arbitrary) scale. If Bragg peaks are modeled as delta functions (Equation 37), the intensity integral is as follows:

$$\begin{aligned} I_{\text{total}} &= \int d^3\mathbf{s}\,[I_D(\mathbf{s}) + I_B(\mathbf{s}) v_c^* \delta(\mathbf{s}-\mathbf{g_h}) + S(\mathbf{s})] \\ &= v_c^* \sum_{\mathbf{h}} I_B(\mathbf{g_h}) + \int d^3\mathbf{s}\,[I_D(\mathbf{s}) + S(\mathbf{s})]. \end{aligned} \tag{59}$$

As discussed in the previous section, the Bragg intensity and the continuous scattering (proportional to $I_D + S$) are measured on an arbitrary scale. To determine the unknown scale factor, the total measured intensity (Equation 59) can be compared with the calculated theoretical intensity (Equation 58). There are two complications, however. The first is that the Bragg peak at $\mathbf{h}=\mathbf{0}$ is unmeasurable. The second is that the limits of integration in Equations 58 and 59 go to infinity, while the measurement ends at some finite maximum scattering vector, $s_{\max}$. To overcome these complications in the theoretical scattering, the $\mathbf{h}=\mathbf{0}$ Bragg intensity is subtracted and the integral truncated at $s_{\max}$, as follows:

$$I_{\text{total, predicted}} = -Z^2 + \int_0^{s_{\max}} 4\pi s^2 ds \sum_n \left(f_n^2(s) + S_n(s)\right), \tag{60}$$

where $Z = \sum_n f_n(0)$ is the number of electrons in the unit cell. This is compared with the integral of the measured data, over the same region of reciprocal space:

$$I_{\text{total, measured}} = v_c^* \sum_{\mathbf{h}\neq 0} I_B(\mathbf{g_h}) + \int_{|s|<s_{\max}} d^3\mathbf{s}\, I_C(\mathbf{s}), \tag{61}$$

where $I_B$ and $I_C$ are the measured Bragg and continuous intensities on an arbitrary scale. The scale factor is calculated from the ratio of predicted and measured total scattering (Equations 60 and 61).

### Appendix 3 Lattice Dynamics Simulation

Lattice dynamics techniques are used to calculate the vibrational frequencies and normal modes (dispersion relations) of atomic or molecular crystals (*38, 39*), often in the context of diffuse scattering (*40, 41*). Here we consider a class of lattice models where the forces between atoms are modeled as a network of springs, and the atoms are assigned to groups that move collectively as rigid bodies.

**Equations of motion in a rigid-body coordinate system.** The equations of motion for the network can be expressed using a generalized coordinate system for rigid-body displacement. The instantaneous displacement of a rigid group from its resting position is described by a six-coordinate vector, $\mathbf{w} = (t_1, t_2, t_3, \lambda_1, \lambda_2, \lambda_3)$ , where $\mathbf{t}$ and $\boldsymbol{\lambda}$ are the vectors of translation and libration (*42*). For small rotations, the displacement $\mathbf{u}$ of an atom in the group is related to $\mathbf{w}$ by a linear operator: $\mathbf{u} = \mathbf{A}(\mathbf{r})\mathbf{w}$, where $\mathbf{A}$ depends on the coordinates of the atom at rest, $\mathbf{r} = (r_1, r_2, r_3)$, as follows:

$$\mathbf{A}(\mathbf{r}) = \begin{bmatrix} 1 & 0 & 0 & 0 & r_3 & -r_2 \\ 0 & 1 & 0 & -r_3 & 0 & r_1 \\ 0 & 0 & 1 & r_2 & -r_1 & 0 \end{bmatrix}. \tag{62}$$

In the derivation below, we introduce a compact notation where the cartesian displacements of all $n$ atoms in a unit cell $l$ are represented by a $3n$-dimensional column vector $\mathbf{u}_{(l)}$, and similarly the generalized coordinates of all $m$ groups in the unit cell are represented by a $6m$-dimensional column vector $\mathbf{w}_{(l)}$. A $3n$-by-$6m$ matrix $\mathbf{A}$ is constructed so that

$$\mathbf{u}_{(l)} = \mathbf{A}\,\mathbf{w}_{(l)}. \tag{63}$$

Using this notation, the harmonic part of the total potential energy is

$$V^{(2)} = \frac{1}{2}\sum_{l,l'} \mathbf{w}_{(l)}^{\mathrm{T}} \mathbf{V}_{(l,l')} \mathbf{w}_{(l')}, \tag{64}$$

where $\mathbf{V}_{(l,l')}$ is the portion of the Hessian matrix (or the force constants matrix) pertaining to the interaction between a unit cell $l$ and $l'$, defined as follows:

$$\mathbf{V}_{(l,l')} = \mathbf{A}^{\mathrm{T}} \left. \frac{\partial^2 V}{\partial \mathbf{u}_{(l)}\, \partial \mathbf{u}_{(l')}^{\mathrm{T}}} \right|_{\mathbf{u}=0} \mathbf{A}, \tag{65}$$

where $V$ is the total potential energy. The total kinetic energy is

$$T = \frac{1}{2}\sum_{l} \dot{\mathbf{w}}_{(l)}^{\mathrm{T}} \mathbf{M} \dot{\mathbf{w}}_{(l)}, \tag{66}$$

where $\mathbf{M}$ is the matrix of generalized masses (which includes the moments of inertia), defined as follows:

$$\mathbf{M} = \mathbf{A}^{\mathrm{T}} \left(\mathrm{diag}(\mathbf{m}) \otimes \mathbf{I}_3\right) \mathbf{A}, \tag{67}$$

where diag($\mathbf{m}$) is a square matrix with the atomic masses along the diagonal, $\mathbf{I}_3$ is the $3 \times 3$ identity matrix, and $\otimes$ is the Kronecker product.

The equations of motion can be derived from the kinetic and potential energies (Equations 64 and 66) using Lagrangian mechanics, with the following result:

$$\mathbf{M}\ddot{\mathbf{w}}_{(l)} = -\sum_{l'} \mathbf{V}_{(l,l')} \mathbf{w}_{(l')}. \tag{68}$$

The generalized mass matrix is positive definite and can be decomposed as $\mathbf{M} = \mathbf{L}\mathbf{L}^{\mathrm{T}}$, where $\mathbf{L}$ is a lower triangular matrix (found by Cholesky decomposition).

**Normal modes.** The Born / von-Karman approach (*38–40*) can be used to solve the equations of motion (Equation 68) by transforming from real space to reciprocal space (**k**-space, where **k** is the wavevector). First, displacement-wave solutions are proposed with the following form:

$$\mathbf{w}^{\pm}_{(l)} = \mathbf{L}^{-\mathrm{T}}\mathbf{e}\exp\left[i\left(\mathbf{k}\cdot\mathbf{r}_l \pm \omega t\right)\right], \tag{69}$$

where $\mathbf{e}$ is a (complex) polarization vector, $\mathbf{r}_l$ is the origin of unit cell $l$, $\omega$ is the angular frequency, and $t$ is time. Substituting these solutions into the equations of motion,

$$\omega^2\mathbf{e} = \left\{\mathbf{L}^{-1}\sum_{l'}\mathbf{V}_{(l,l')}\exp\left[i\mathbf{k}\cdot\left(\mathbf{r}_{l'}-\mathbf{r}_l\right)\right]\mathbf{L}^{-\mathrm{T}}\right\}\mathbf{e}. \tag{70}$$

This can be cast in the form of an eigenvalue equation, $\omega^2\mathbf{e} = \mathbf{D}_{(\mathbf{k})}\mathbf{e}$, where $\mathbf{D}_{(\mathbf{k})}$ is known as the dynamical matrix. Let $\omega^2_{(\mathbf{k},j)}$ and $\mathbf{e}_{(\mathbf{k},j)}$ be the $j^{\mathrm{th}}$ eigenvalue and eigenvector of $\mathbf{D}_{(\mathbf{k})}$. These satisfy

$$\omega^2_{(\mathbf{k},j)}\mathbf{e}_{(\mathbf{k},j)} = \mathbf{D}_{(\mathbf{k})}\mathbf{e}_{(\mathbf{k},j)}. \tag{71}$$

Here we consider finite supercells consisting of $N$ primitive unit cells and periodic boundary conditions so that $\mathbf{k}$ is restricted to a discrete set. It can be shown (*39*) that the general solution is a sum of all allowed waves, or normal modes, as follows:

$$\mathbf{w}_{(l)} = \sum_{\mathbf{k}}\sum_{j}\sigma_{(\mathbf{k},j)}\mathbf{L}^{-\mathrm{T}}\mathbf{e}_{(\mathbf{k},j)}\exp\left(i\mathbf{k}\cdot\mathbf{r}_l\right), \tag{72}$$

where $\sigma_{(\mathbf{k},j)}$ are complex, time-dependent amplitudes of the normal modes.

**Displacement covariances for thermally-excited vibrations.** In thermodynamic equilibrium, the mean-squared amplitudes of the normal coordinates can be found by applying the equipartition theorem (high-temperature limit):

$$\left\langle\left|\sigma_{(\mathbf{k},j)}\right|^2\right\rangle = N^{-1}k_BT\omega^{-2}_{(\mathbf{k},j)}. \tag{73}$$

Using this result, the covariance matrix can be derived for the generalized coordinates. It can be expressed in the following compact form:

$$\left\langle\mathbf{w}_{(l)}\mathbf{w}^{\mathrm{T}}_{(l')}\right\rangle = N^{-1}k_BT\sum_{\mathbf{k}}\exp\left(i\mathbf{k}\cdot\left(\mathbf{r}_l-\mathbf{r}_{l'}\right)\right)\mathbf{K}^{+}_{(\mathbf{k})}, \tag{74}$$

where $\mathbf{K}_{(\mathbf{k})}$ is the Fourier transform of the force constants matrix ($\mathbf{K}_{(\mathbf{k})} = \mathbf{L}\mathbf{D}_{(\mathbf{k})}\mathbf{L}^T$) and $\mathbf{K}^{+}_{(\mathbf{k})}$ is its generalized inverse (in the generalized inverse, normal modes corresponding to rotation and translations of the entire crystal have $\omega = 0$ and are excluded). The covariance of individual atomic displacements can be calculated by projection (Equation 62), as follows:

$$\left\langle\mathbf{u}_{(jl)}\mathbf{u}^{\mathrm{T}}_{(j'l')}\right\rangle = \mathbf{A}(\mathbf{r}_j)\left\langle\mathbf{w}_{(\kappa l)}\mathbf{w}^{\mathrm{T}}_{(\kappa' l')}\right\rangle\mathbf{A}(\mathbf{r}_j)^{\mathrm{T}}, \tag{75}$$

where the index $\kappa$ refers to the rigid group to which atom $j$ belongs ($\mathbf{w}_{(\kappa l)}$ is a 6-element vector). The self-terms of the covariance matrix are the familiar atomic displacement parameters (ADPs) in protein crystallography:

$$\mathbf{U}_j = \left\langle\mathbf{u}_{(j)}\mathbf{u}^{\mathrm{T}}_{(j)}\right\rangle. \tag{76}$$

The equivalent isotropic B-factor is defined as

$$B_{\mathrm{eq}} = \left(8\pi^2/3\right)\mathrm{tr}\left(\mathbf{U}\right). \tag{77}$$

Because the generalized coordinates of the lattice model $\mathbf{w}$ are the same as those used in TLS refinement (*42*), the $\mathbf{T}$, $\mathbf{L}$, and $\mathbf{S}$ matrices are $3\times 3$ blocks within the larger covariance matrix for $\mathbf{w}$. For a particular rigid group $\kappa$, the matrices are related as follows:

$$\left\langle\mathbf{w}_{(\kappa)}\mathbf{w}^{\mathrm{T}}_{(\kappa)}\right\rangle = \begin{bmatrix}\mathbf{T} & \mathbf{S}^{\mathrm{T}}\\ \mathbf{S} & \mathbf{L}\end{bmatrix}. \tag{78}$$

**Diffuse scattering and the one-phonon approximation.** It is customary in phonon scattering theory to represent the reciprocal space coordinate by the momentum transfer vector $\mathbf{q}$ rather than the scattering vector $\mathbf{s}$, with the relationship $\mathbf{q} = 2\pi\mathbf{s}$. We follow this convention below. Since the lattice vibrations are harmonic, the diffuse scattering intensity per unit cell can be calculated in the harmonic approximation:

$$I_D = N^{-1} \sum_{l,l'} e^{i\mathbf{q}\cdot(\mathbf{r}_l - \mathbf{r}_{l'})} \sum_{j,j'} f_j f_{j'} e^{i\mathbf{q}\cdot(\mathbf{r}_j - \mathbf{r}_{j'})} T_j T_{j'} \left[T_{lj,l'j'} - 1\right]. \tag{79}$$

The first summation is over all pairs of unit cells indexed by $l$ and $l'$, and $\mathbf{r}_l$ is the origin of unit cell $l$. The second summation is over all atom pairs in unit cell $l$ (indexed by $j$) and $l'$ (indexed by $j'$). Here, $f_j$ is the atomic scattering factor and $\mathbf{r}_j$ is the average position of the atom relative to the unit cell origin. The factors $T_j$ (the Debye-Waller factor) and $T_{lj,l'j'}$ depend on the self- and cross-terms of the displacement covariance matrix, as follows:

$$T_j = \exp\left[-\frac{1}{2}\mathbf{q}^{\mathrm{T}}\mathbf{U}_j\mathbf{q}\right] \tag{80}$$

and

$$T_{lj,l'j'} = \exp\left[\frac{1}{2}\mathbf{q}^{\mathrm{T}}\left(\langle\mathbf{u}_{j'l'}\mathbf{u}_{jl}^{\mathrm{T}}\rangle + \langle\mathbf{u}_{jl}\mathbf{u}_{j'l'}^{\mathrm{T}}\rangle\right)\mathbf{q}\right], \tag{81}$$

where $\mathbf{u}_{jl}$ is the instantaneous displacement of atom $j$ in unit cell $l$ and $\mathbf{U}_j$ is defined in Equation 76.

When the cross-terms of the covariance matrix are small, Equation 81 can be expanded as a Taylor series, and the intensity separated into terms of increasing order ($I = I^{(1)} + I^{(2)} + \ldots$) where $I^{(1)}$ is the "one-phonon" diffuse scattering, $I^{(2)}$ is the "two-phonon" diffuse scattering, and so on. In the "one-phonon" term, a phonon with wavevector $\mathbf{k}$ contributes only to points in reciprocal space that are displaced by $\mathbf{k}$ from the Bragg peak locations $\mathbf{g_h}$, as follows:

$$I^{(1)}(\mathbf{q} = \mathbf{g_h} - \mathbf{k}) = \sum_n \frac{k_B T}{\omega^2_{(\mathbf{k},n)}} \left| \sum_j f_j T_j \exp\left[i\mathbf{q}\cdot\mathbf{r}_j\right] \mathbf{q}^{\mathrm{T}}\mathbf{A}(\mathbf{r}_j)\mathbf{L}^{-\mathrm{T}}_{(\kappa)}\mathbf{e}_{(\kappa,\mathbf{k},n)} \right|^2. \tag{82}$$

This can be written in a more compact form that is computationally convenient by defining the vector-valued "one-phonon structure factor",

$$\mathbf{G}(\kappa, \mathbf{q}) = \mathbf{q}^{\mathrm{T}} \sum_j f_j T_j \exp\left[i\mathbf{q}\cdot\mathbf{r}_j\right] \mathbf{A}(\mathbf{r}_j). \tag{83}$$

Then, the one-phonon intensity can be written as follows:

$$I^{(1)}(\mathbf{q} = \mathbf{g_h} - \mathbf{k}) = k_B T\, \mathbf{G}(\mathbf{q})\mathbf{K}^{+}_{(\mathbf{k})}\mathbf{G}^{\dagger}(\mathbf{q}), \tag{84}$$

where $\mathbf{K}^{+}_{(\mathbf{k})}$ is the generalized inverse of the force constants matrix (see Equation 74).

### Supplementary Figures

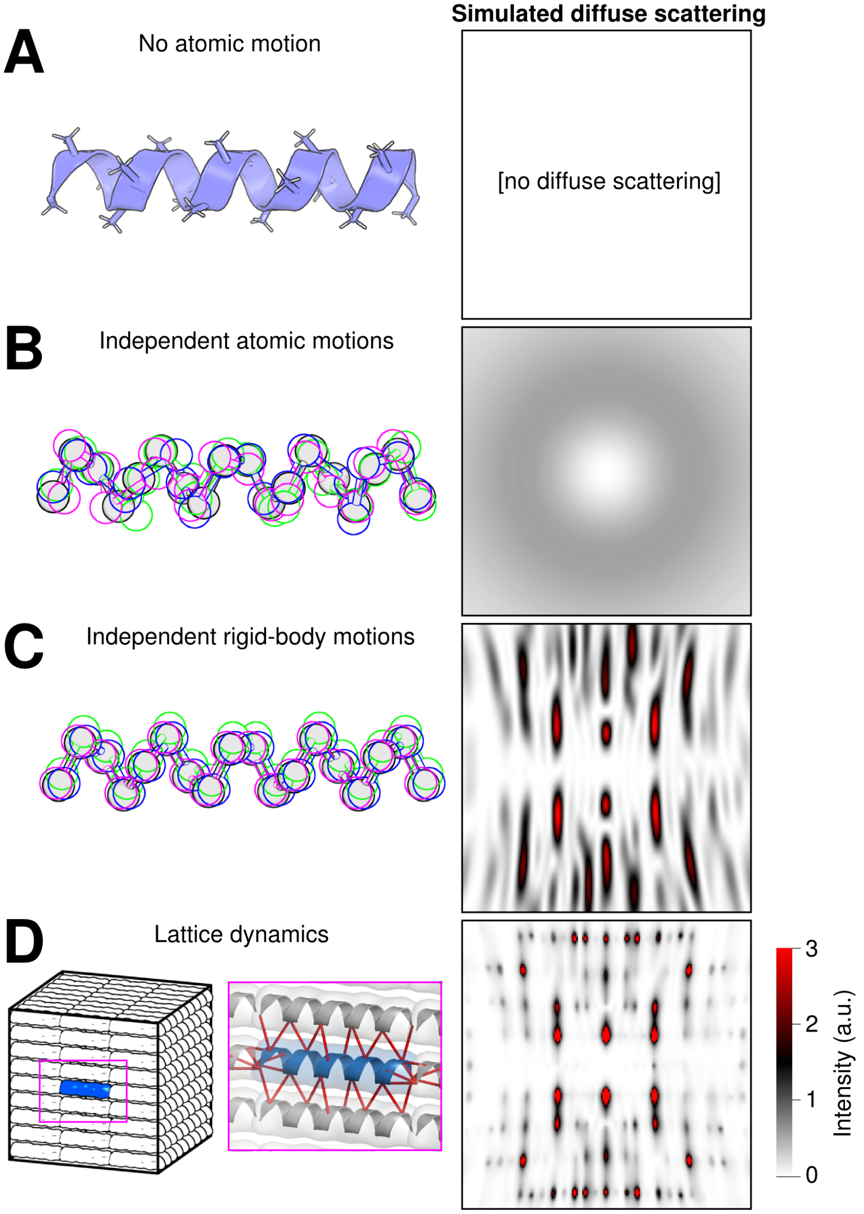


**Fig. S1:** Diffuse scattering depends on how atomic displacements are correlated. To illustrate, scattering patterns were simulated for a crystal of an idealized alpha helix (panel A) with various types of motion. The simulated X-ray beam had a wavelength of 1 Å and was perpendicular to the helical axis (into the page). (A) If there is no atomic motion (left), then there is no diffuse scattering (right). (B) Uncorrelated atomic displacements with a B-factor of 20 $\text{Å}^2$. On the left, a representative ensemble is illustrated for $C_\alpha$ positions with the mean shaded in gray and three random samples outlined in different colors. The diffuse scattering is relatively featureless and isotropic (right). (C) Translational rigid body motion that is uncorrelated between unit cells. B-factors are the same as in panel B. The diffuse scattering (right) has oscillations related to the molecular transform (*43*). (D) Helices move as in panel C except that motions are correlated between unit cells. The correlations were calculated from a vibrational lattice dynamics model, similar to the one developed for this study (Supplementary Methods), consisting of a periodic supercell (left) with a network of springs (red lines, inset). The potential energy function was chosen (Equation 5) so that thermally excited vibrations at room temperature produced the same B-factors as in panels B-C. The simulated diffuse pattern (right) contains intense halo features that coincide with the Bragg peaks (not shown). Although the models in panels B-D have identical B-factors and cannot be distinguished by Bragg diffraction, they predict very different diffuse scattering.

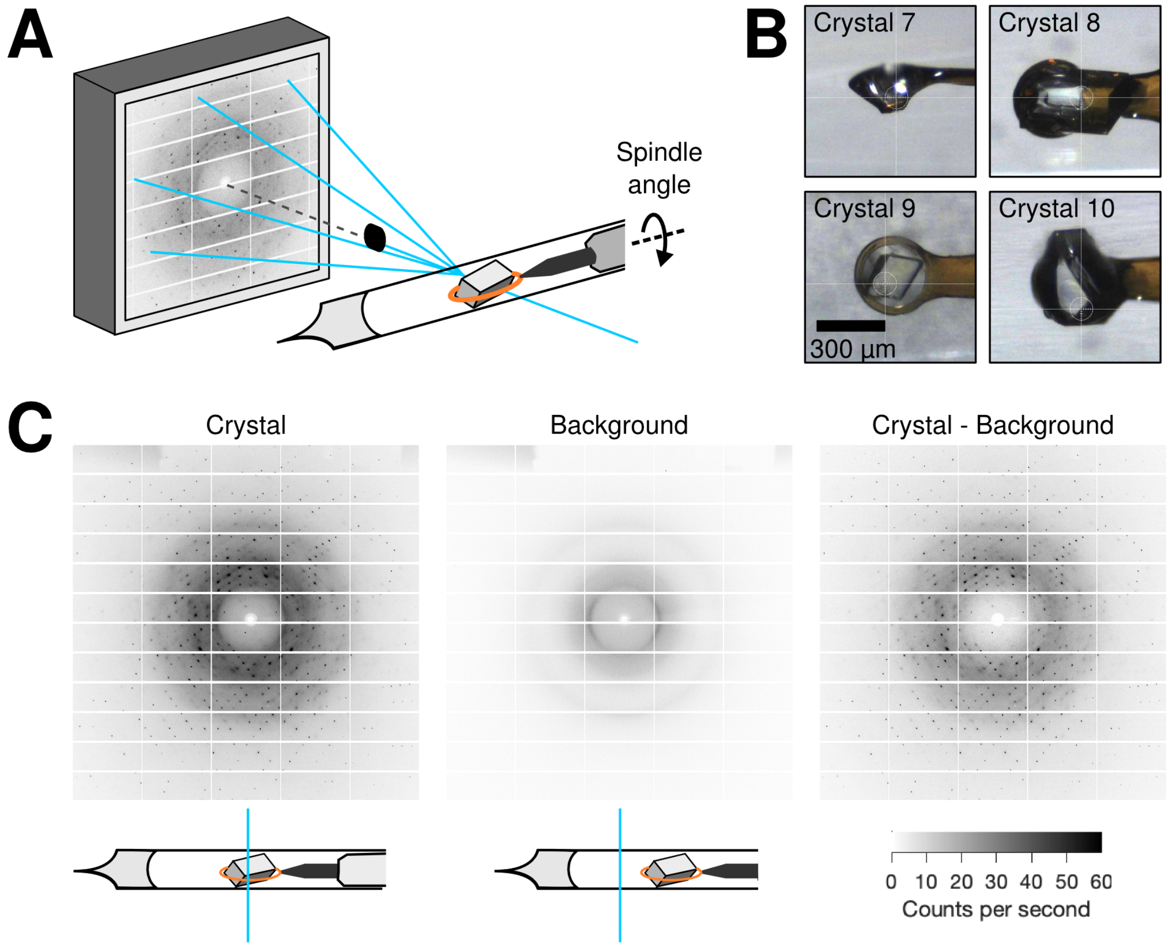


**Fig. S2:** X-ray data collection strategy. (A) Diagram of the experimental setup. To prevent dehydration at ambient temperature, each crystal was enclosed in a plastic capillary with crystallization solution in the tip. X-ray diffraction images were collected using a photon-counting detector using fine phi-slicing (0.1° per frame). (B) The total dose was distributed over 11 diffraction volumes from four large crystals (Table S1). (C) At each spindle angle (phi), diffraction images were collected with the crystal in the beam (left) and with the crystal translated out of the beam (middle). As the background includes scattering from the instrument and narrow rings due to the plastic capillary, the pattern due to the sample only is estimated by subtraction (right). For clear illustration, the images shown are the sums of 10 sequential frames (1° oscillation and 1 s exposure in total).

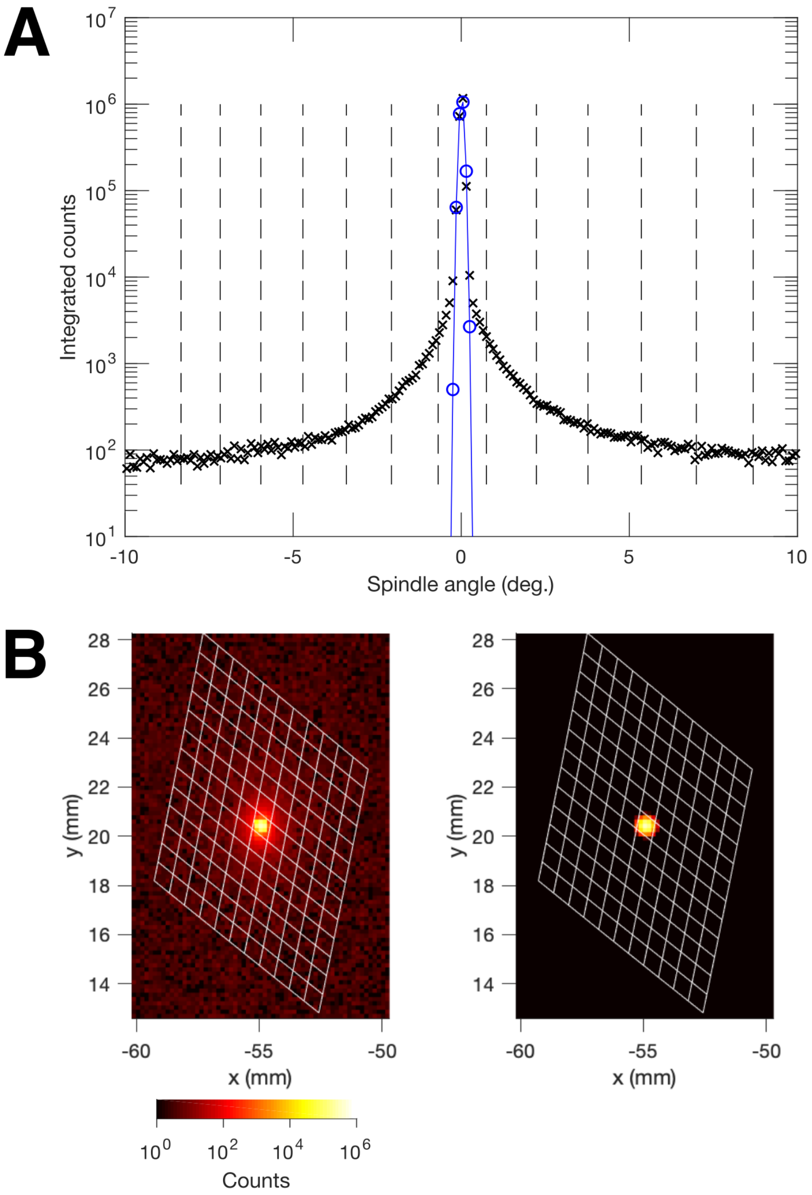


**Fig. S3:** Rocking curve for the (2,-8,-1) reflection. In our diffraction data from triclinic lysozyme, the apparent crystal mosaicity of 0.02 to 0.03° is smaller than the oscillation width of 0.1° per frame, so most Bragg reflections appear only in one or two consecutive images and their rocking curves cannot be resolved. We thus identified an intense (2,-8,-1) Bragg reflection that is close to the spindle axis and passes through the Ewald sphere along $\mathbf{c}^*$ at a rate approximately 3 times slower than the fastest route. The peak profile above the background was well-fit by a three-dimensional Gaussian with apparent mosaicity and beam divergence of 0.020° and 0.024°, respectively. (A) The total photon counts in a 25-pixel region around the Bragg peak is plotted vs. spindle angle from the diffraction maximum (black symbols). The sharp Bragg peak (Gaussian fit shown in blue) is easily distinguished from the more slowly-varying background. The vertical dashed lines show the voxel subdivisions along $\mathbf{c}^*$ used in the construction of the fine diffuse map. The peak is confined to the central subdivision. (B) On the left is a zoom-in of the (2,-8,-1) Bragg reflection in the diffraction image (0.1° oscillation) at its peak intensity. The x and y axes are distances from the beam center, and the spindle is parallel to the x-axis. On the right is the same oscillation simulated using the Gaussian fit in panel A. In both images, the intersection of voxel edges in the $\mathbf{a}^*$-$\mathbf{b}^*$ plane with the Ewald sphere are projected onto the detector plane, showing that the Bragg intensity is confined to the central voxel.

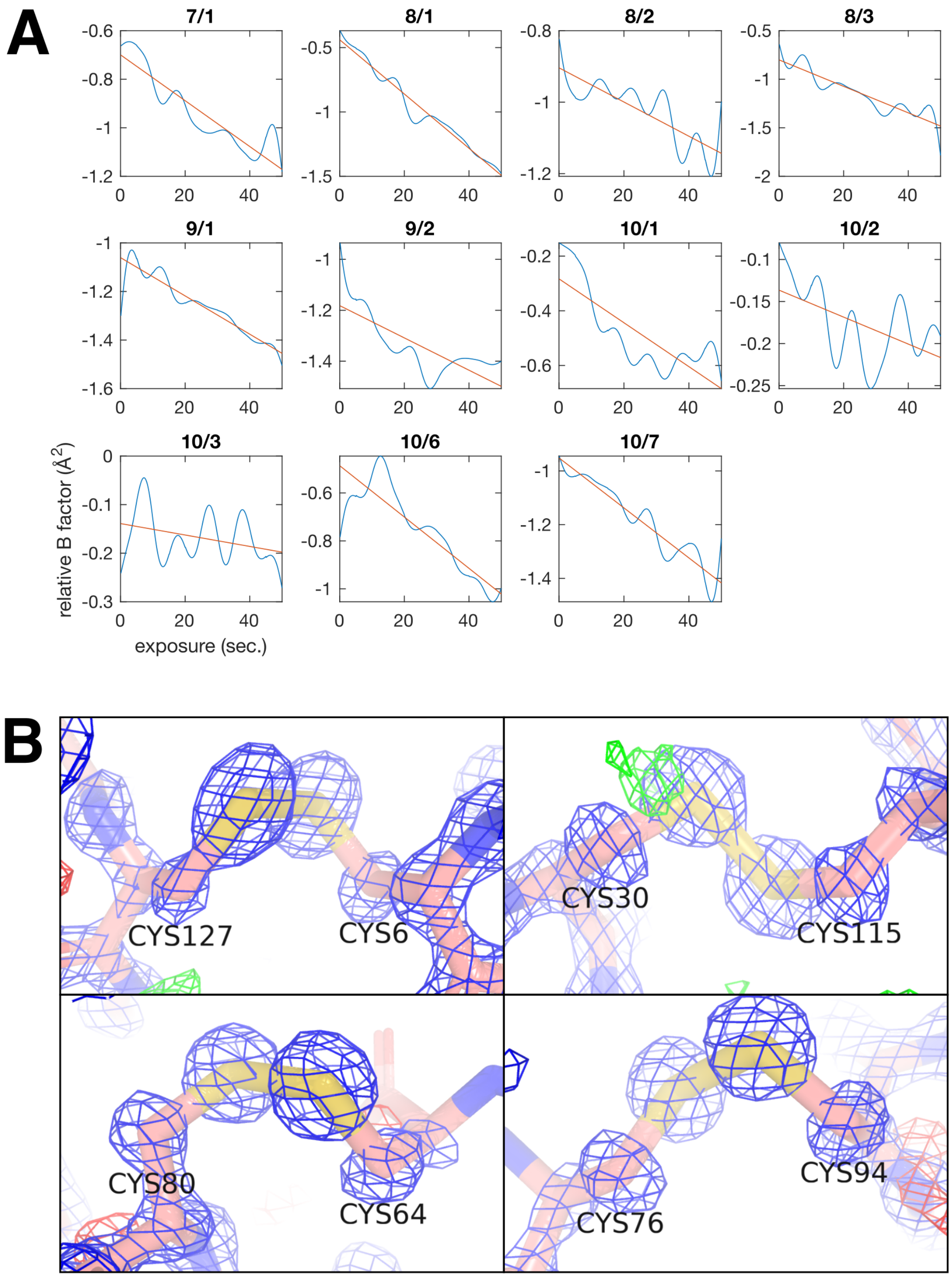


**Fig. S4:** Minimal radiation damage is observed in the triclinic lysozyme dataset. (A) In general, global radiation-induced damage manifests as a decay in the Wilson B-factor with increasing dose, which is corrected by a relative B-factor when scaling Bragg data with *aimless* (*5*). The relative B-factor (blue curves) and a linear fit (red lines) are shown for all 11 diffraction volumes from four crystals (Fig. S2B) with labels corresponding to crystal number / wedge number (Table S1). For each of the wedges used here, the Wilson B-factor decays by less than 1 Å$^2$ over the course of the measurement. (B) Disulfide bonds were also inspected for evidence of photo-reduction in the structure determined from the combined data. The four disulfide bonds in the lysozyme are shown along with the 2Fo-Fc map (blue mesh at 2.5$\sigma$) and the Fo-Fc map (red and green mesh at $\pm$2.5$\sigma$). The bonds appear largely intact. The difference density near the sulfur atom of C30 (top right panel) may indicate that a minor population of C30-C115 has been reduced.

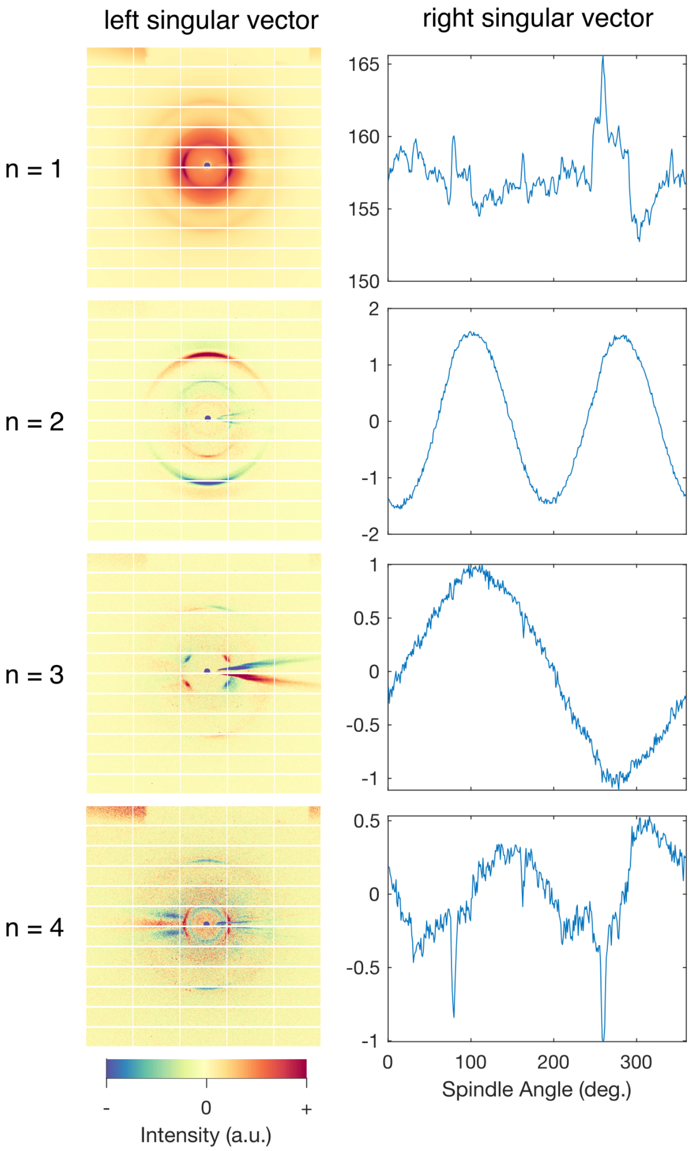


**Fig. S5:** Importance of background subtraction. Singular value decomposition of the set of background images (360°, 1 s exposures, 1° oscillation) shows several scattering features that vary independently as the spindle rotates. The first four singular vectors are shown ($n = 1$ to 4); the left singular vectors correspond to diffraction images, and the right singular vectors show the contribution as a function of spindle angle. The $n = 1$ component contains the overall background scattering. The $n = 2$ component, which has ring-like features and a period of 180°, can be attributed to anisotropic scattering of the capillary. The $n = 3$ component, with strong equatorial features and a period of 360°, is due to the shadow cast by the sample pin as it precesses. The remaining components ($n > 3$) have features due to both the pin shadow and the capillary, as well as experimental noise.

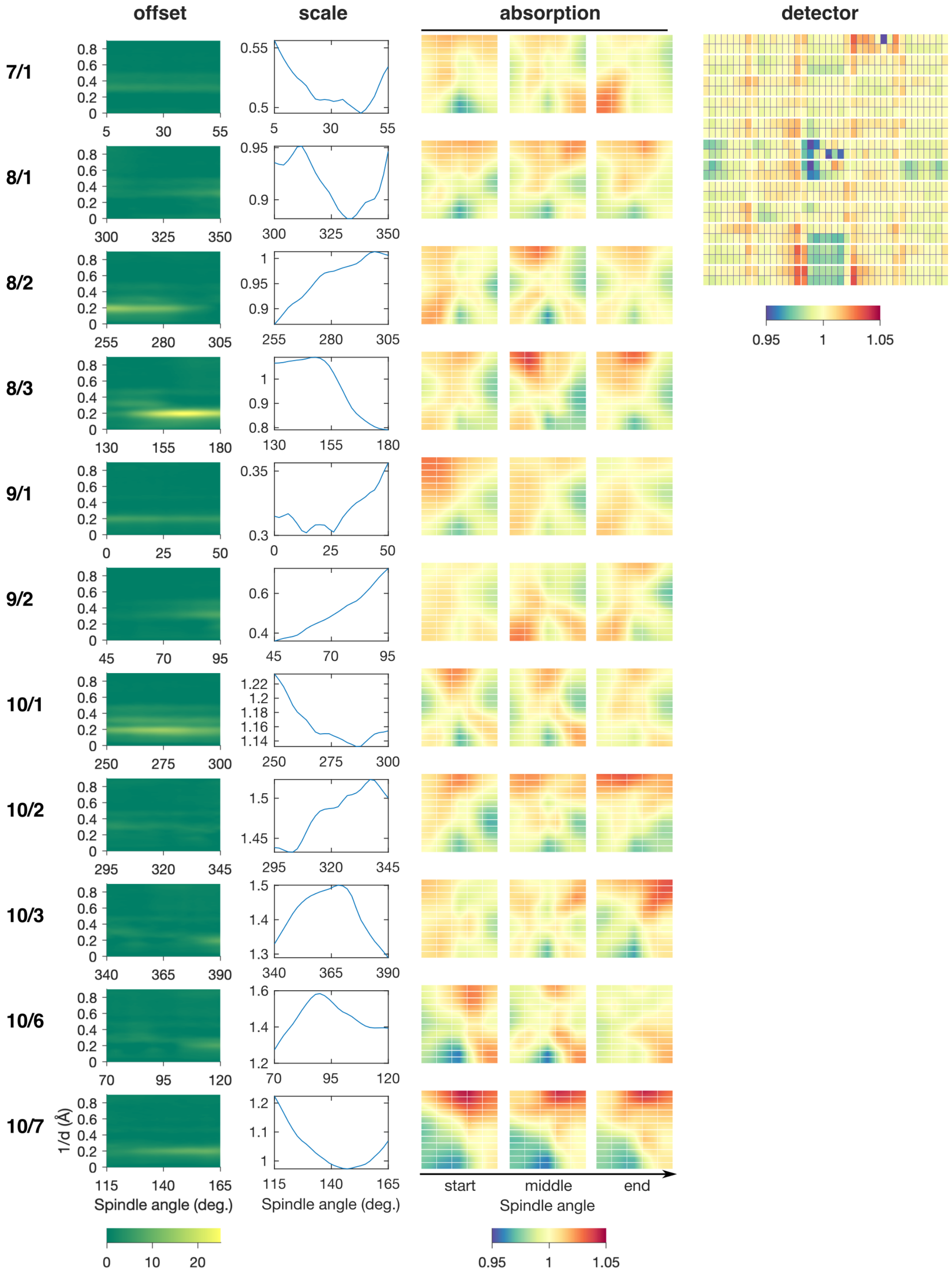


**Fig. S6:** Refinement of a scaling model to correct for experimental artifacts in X-ray images. The scaling model (Equation 1 in Supplementary Methods) used in processing diffuse scattering data relates the merged intensity to the observed intensity as a function of spindle angle, resolution ($d$), and detector position. From left to right, the model parameters included: (1) offset correction vs. spindle angle and resolution, (2) overall scale factor vs. spindle angle, (3) scale factor for absorption correction vs. spindle angle and detector position, and (4) scale factor for detector efficiency correction vs. detector chip index. The parameters in the model were refined in order to minimize the least-squares error between observation and prediction (Equation 2 in Supplementary Methods). The best-fit-parameters are illustrated for each partial dataset (top to bottom) labeled according to crystal number / wedge number (refer to Fig. S2B, Table S1).

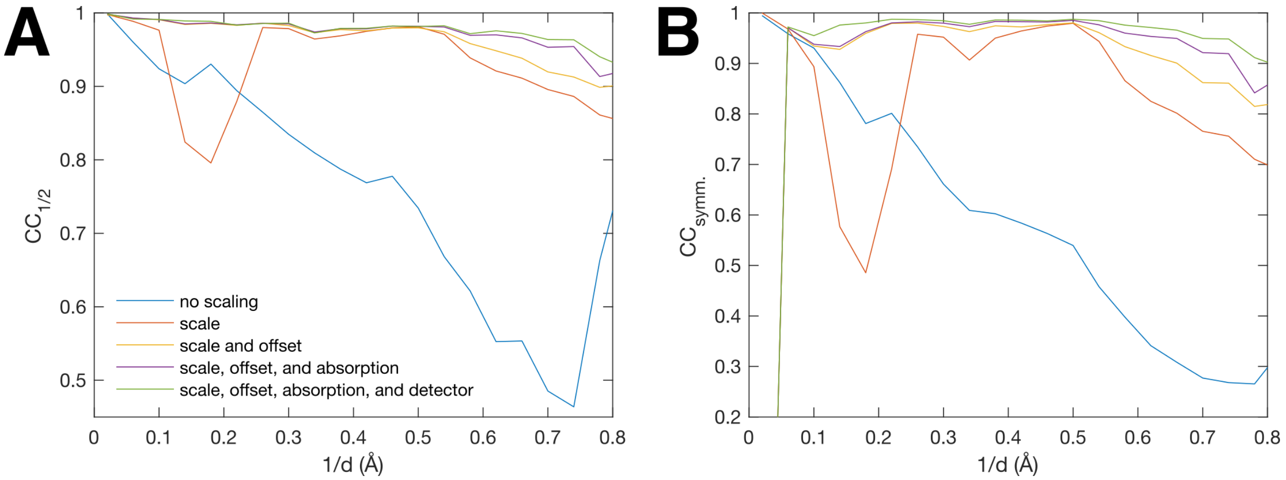


**Fig. S7:** Impact of scaling parameters on diffuse data quality. Four parameters were refined sequentially to the coarse continuous scattering map (one point per integer h,k,l): scale, offset, absorption, and detector (labeled in Fig. S6 and described in Supplementary Methods). Data quality was assessed by comparing Pearson correlation coefficients (CC) between half-datasets within shells of constant resolution. (A) CC between random half-datasets ($CC_{1/2}$). (B) CC between half-datasets related by Friedel symmetry ($l > 0$ and $l < 0$, $CC_{symm.}$). Both metrics show improvements in data quality as each parameter is added.

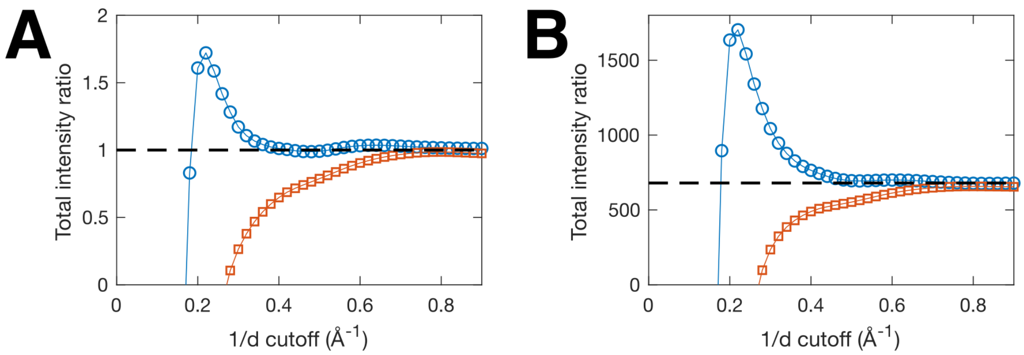


**Fig. S8:** Robustness of absolute intensity calibrations performed with the Krogh-Moe (K-M) method (red squares) and the modified K-M method, which accounts for an interference term that depends on bond geometries (blue circles). (A) A test dataset of X-ray intensities (Bragg, diffuse, and Compton) was calculated from the 1-unit cell MD simulation of triclinic lysozyme, and the ratio between predicted and actual total intensity was plotted vs. resolution cutoff using each method. Both converge to the expected value of 1 (dashed line), but the modified method converges much more quickly. (B) The same methods applied to the experimental maps of continuous scattering and Bragg diffraction on an arbitrary intensity scale. The modified K-M method converged by $\sim 2$ Å resolution, yielding a scale factor of 680 (dashed line).

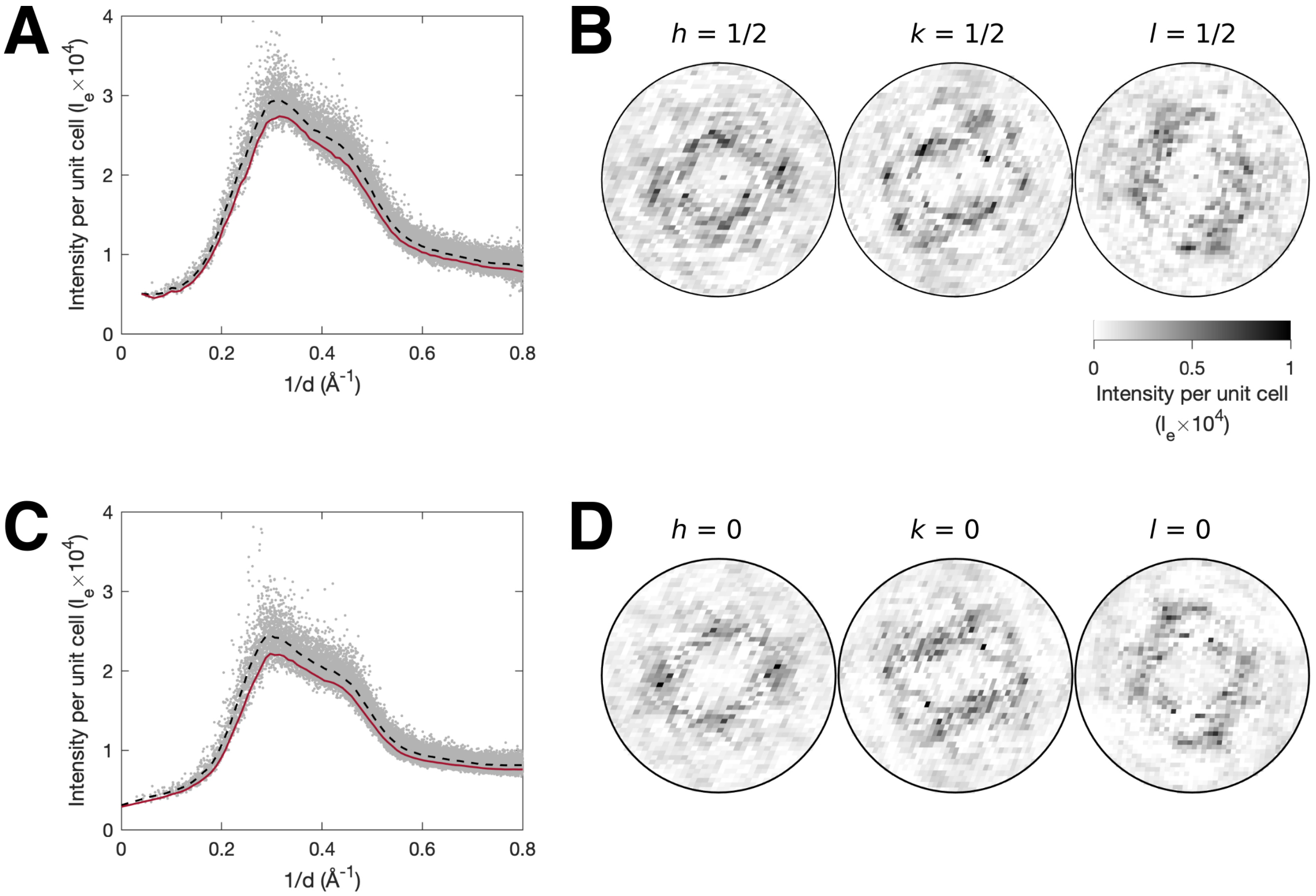


**Fig. S9:** Isotropic component of diffuse scattering in experiment and the 1-unit cell MD simulation. (A) The diffuse scattering was resampled at half-integer values of h, k, and l, where halo scattering is minimal. To improve the signal-to-noise ratio, the interpolation was performed by least-squares fitting a 2nd order polynomial to the 5x5x5 voxel region around each point in the target grid. The resulting "half-integer map" (gray points) was analyzed as a function of the resolution $d$. Within each resolution bin, the isotropic background (red curve) was defined as $1\sigma$ below the mean (dashed black curve). (B) The isotropic background was subtracted from the total intensity to aid in visualizing the cloudy variational pattern. Slices from left to right are perpendicular to $\mathbf{a}^*$, $\mathbf{b}^*$, and $\mathbf{c}^*$. (C-D) The 1-unit cell MD simulation was analyzed in the same way as the half-integer map. The isotropic scattering and variational scattering components are similar between the 1-unit cell MD simulation and the half-integer map.

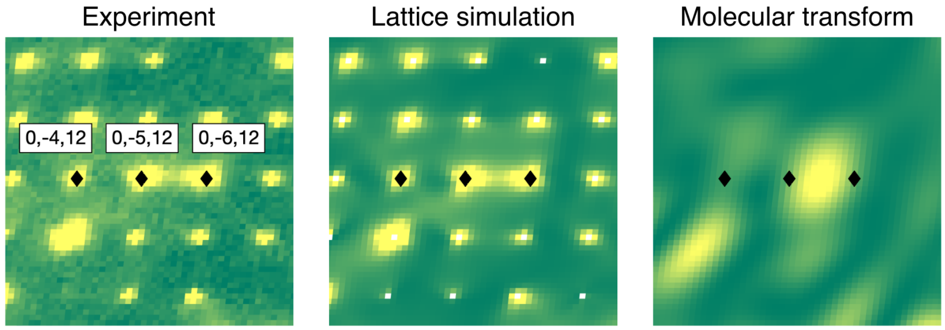


**Fig. S10:** Diffuse streaks attributed to intense features in the molecular transform. In the experimental maps of variational scattering (left panel), certain pairs of halos are connected by streaks of diffuse intensity, such as (0,-5,12) and (0,-6,12), while others are not, such as (0,-4,12) and (0,-5,12). Similar features are seen in the one-phonon scattering simulation of the lattice dynamics model (center panel). These anisotropic features occur because the halo intensities are modulated by the molecular transform, the continuous diffraction intensity of an isolated unit cell (right); the molecular transform is intense between (0,-5,12) and (0,-6,12) but is close to zero between (0,-4,12) and (0,-5,12), explaining the presence of a diffuse streak in the former case and its absence in the latter.

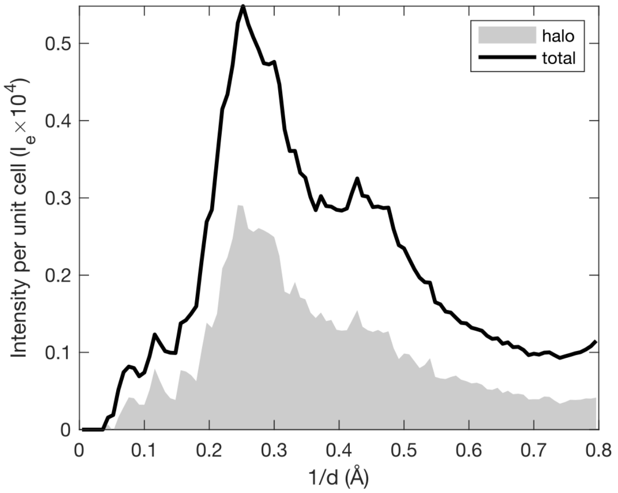


**Fig. S11:** Contribution of halos to the total variational scattering. The mean variational intensity in each resolution shell was calculated for the full map, which includes the halo features, and for the half-integer interpolated map (described in Fig. S9A)t. The halo contribution was estimated as the difference between the two (shaded region). In most resolution shells, the halos account for about half of the total variational intensity (solid curve).

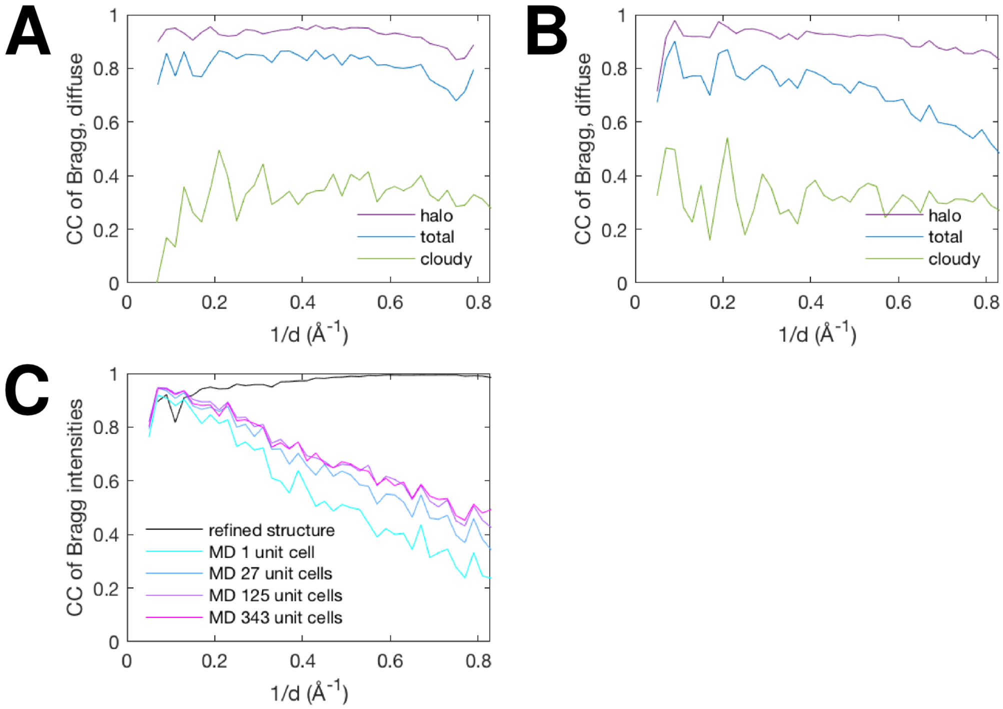


**Fig. S12:** Correlations between Bragg and diffuse intensities. The total variational intensity, which includes halos and cloudy scattering, was averaged in neighborhoods ($h$,$k$,$l$) around each Bragg peak ($h_0, k_0, l_0$), where $|h - h_0| < 1/2$, $|k - k_0| < 1/2$, and $|l - l_0| < 1/2$. The contribution of cloudy scattering was estimated by linear interpolation of the half-integer map (as was done in Fig. S9A). Halo scattering was estimated by subtracting the cloudy scattering from the total variational intensity. (A) The Pearson correlation coefficients (CC) were calculated between the Bragg intensities and different components of diffuse scattering: total variational scattering (blue curve), cloudy scattering (green curve), and halo scattering (purple curve). The correlations are positive in all resolution shells, with halo intensities showing the strongest correlation. (B) The diffuse scattering and Bragg intensities calculated from the 343-unit cell MD simulation were analyzed as in panel A. The Bragg intensities and halo intensities are also strongly correlated. (C) CC of Bragg intensities between experiment and four MD simulations with 1-343 unit cells (colored curves, see legend). The CC is strong at low resolution, but decays significantly at high resolution. In contrast, the CC between the experiment and the refined crystal structure model (black curve) remains excellent at high resolution.

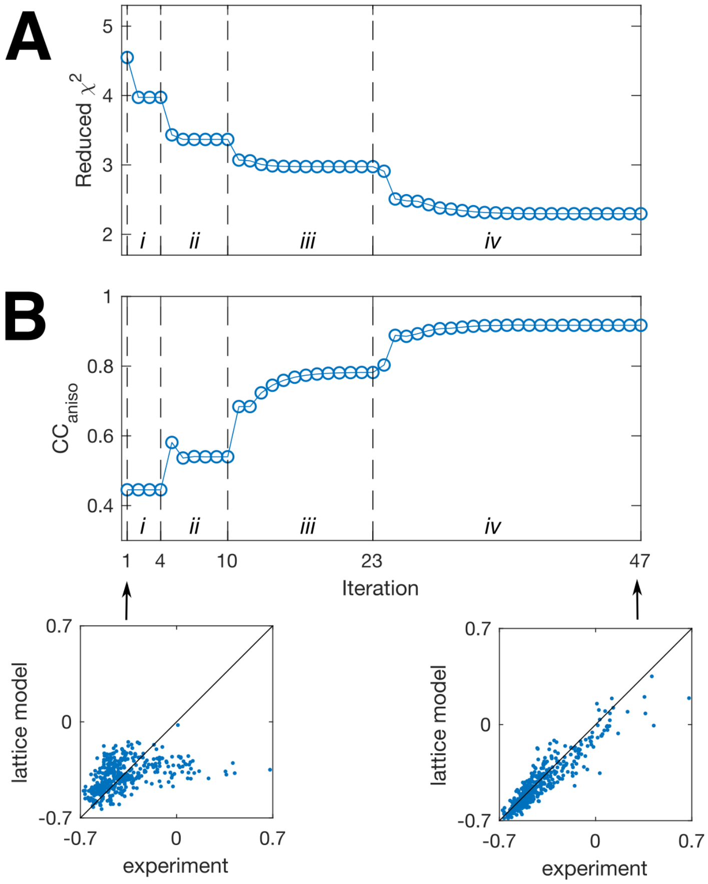


**Fig. S13:** Refinement of lattice dynamics model against a small subset of diffuse halos. (A) The spring constants in the lattice dynamics model were refined iteratively to minimize the least-squares difference (reduced $\chi^2$, blue symbols) between the calculated one-phonon intensity and the measured variational intensity around 400 intense halos. To avoid over-fitting, the refinement was carried out in four stages (i-iv) as restraints were removed (Supplementary Methods). The first stage (i) is a Gaussian model with equal springs, while in the final stage (iv) all springs are refined individually. (B) The ability of the model to explain halo anisotropy (see Main Text and Fig. 3C) was assessed using the Pearson correlation coefficient for the 400 halos ($CC_{aniso}$). Initially, the anisotropy parameters for model and experiment show little correlation ($CC_{aniso} = 0.45$, scatter plot in lower left). After refinement, the correlation improves ($CC_{aniso} = 0.92$, scatter plot in lower right).

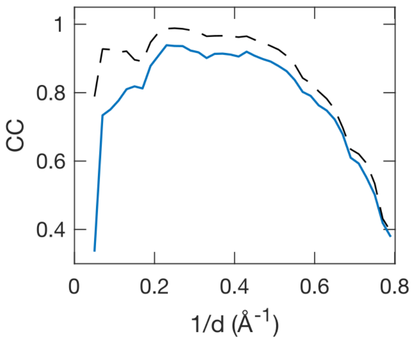


**Fig. S14:** Correlation of experiment and simulated one-phonon scattering from the refined lattice dynamics model calculated with intensities from the full 13x11x11 grid. The data-model correlation (solid blue curve) is excellent (CC $\sim$ 0.9) between 2 and 5 Å resolution (solid), but decays rapidly beyond $\sim$ 2 Å resolution. At high resolution, the correlation is limited by noise in the experimental map, as estimated using the CC* statistic (dashed curve). Interpolating the maps on a 7x7x7 grid improves the signal-to-noise (Fig. 2D).

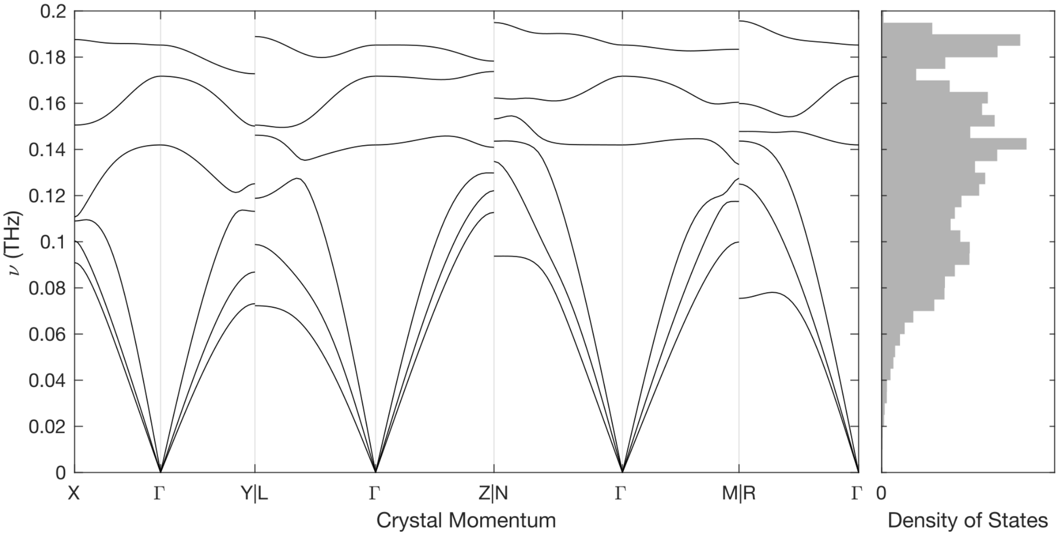


| | $b_1$ | $b_2$ | $b_3$ |
|---|---|---|---|
| Γ | 0 | 0 | 0 |
| L | 0 | 0.5 | -0.5 |
| M | 0.5 | 0 | 0 |
| N | 0.5 | -0.5 | -0.5 |
| R | 0.5 | 0 | -0.5 |
| X | 0 | 0 | -0.5 |
| Y | 0 | 0.5 | 0 |
| Z | 0.5 | -0.5 | 0 |

**Fig. S15:** Dispersion relations for lattice modes of triclinic lysozyme. The vibrational frequencies ($\nu = 2\pi\omega$) were calculated from the refined lattice dynamics model along a path through k-space (x-axis on the left plot) that passes through the origin ($\Gamma$) as well as special points at the edge of the Brillouin zone (defined in the table on the right, where $b_1$, $b_2$, and $b_3$ are the fractional coordinates along a*, b*, and c*). The vibrational density of states (right plot) was calculated for 11,025 evenly spaced points in the Brillouin zone (a 25x21x21 supercell). In the model, each protein chain has 6 degrees of freedom (3 coordinates each for translations and rotations), resulting in 6 vibrational modes: 3 optical and 3 acoustic. The acoustic modes are those that pass through the origin.

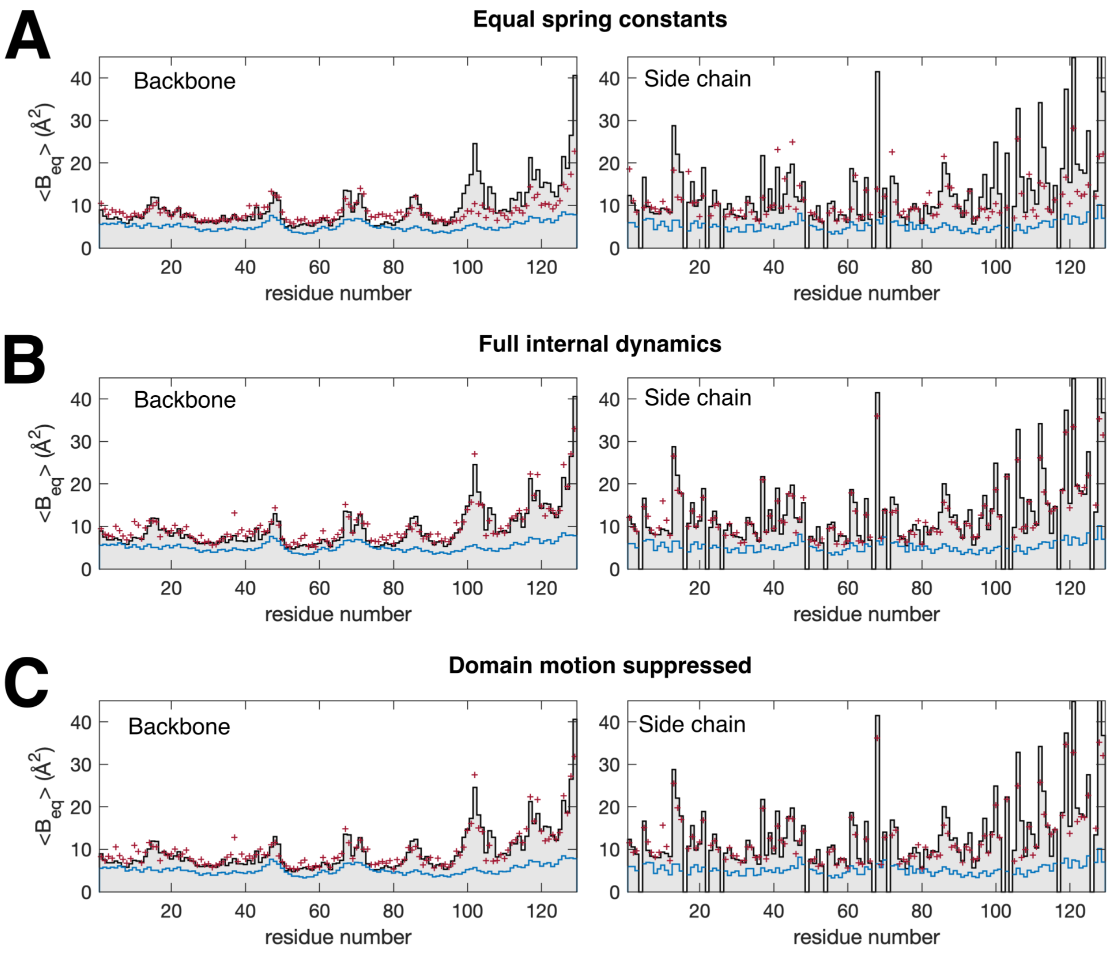


**Fig. S16:** Elastic network models of the internal motion refined to the total atomic displacement parameters (ADPs). For visual clarity, the average equivalent isotropic B-factors were computed for non-H atoms in the backbone and side-chains for the refined crystal structure (black curve and gray shaded area), the lattice dynamics model (blue curve), and the lattice + internal dynamics models (dark red symbols). (A) A model with equal spring constants reproduces many of the backbone features, except at the C-terminus (residues 95-129). Many of the side-chain B-factors are not reproduced. (B) The B-factors are reproduced by refining the spring constants in a model parameterized with one coupling constant per residue (see Fig. 4C). Agreement for the backbone (left) and side-chain (right) B-factors is improved. (C) Repeating the refinement in panel B with a modified potential energy function, which suppresses collective motion of the $\alpha$ and $\beta$ domains (see Fig. 4D, Supplementary Methods), lead to a similarly reasonable fit to the B-factors.

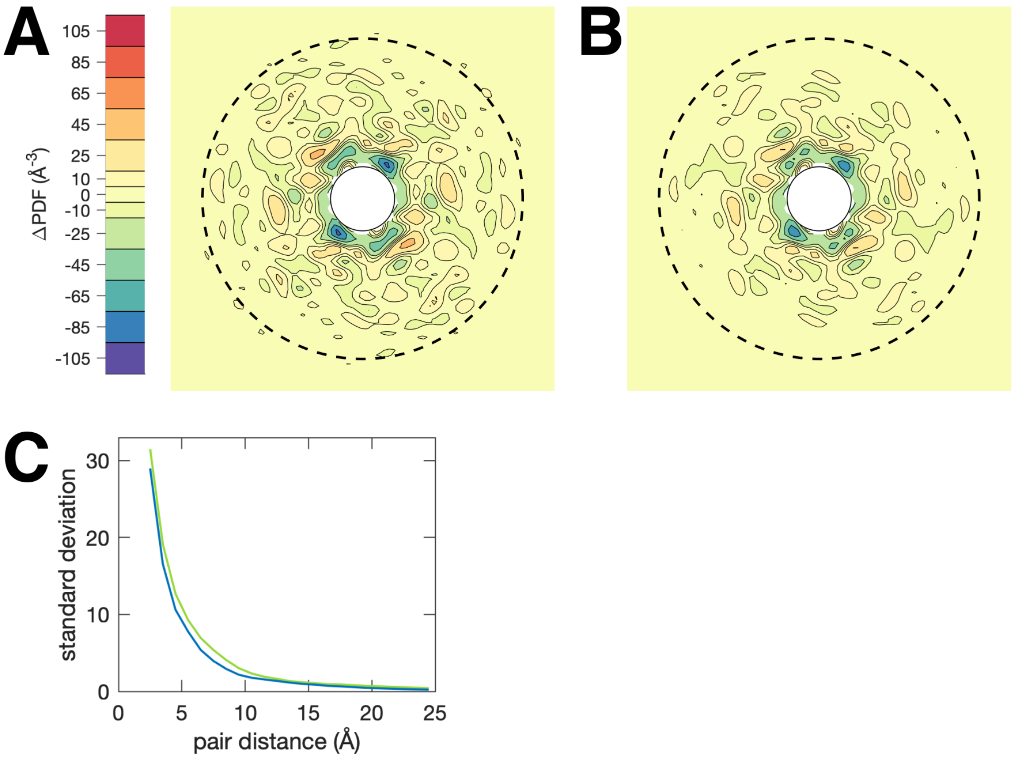


**Fig. S17:** Comparison of models for internal motion by diffuse Patterson analysis. The central sections of the diffuse Patterson maps in the in **a-b** plane (dashed circle corresponds to 10 Å) are shown for the two elastic network models with fit equivalently well to the experimental ADPs (Fig. S16B-C). (A) Fluctuations are observed to $\sim 10$ Å when full internal motions are allowed (figure panel repeated from Fig. 5C). (B) Suppressing collective motion of the $\alpha$ and $\beta$ domains in lysozyme leads to a diffuse Patterson map that is significantly altered. (C) The amplitudes of the fluctuations in the diffuse Patterson for the no-domain-motion model (blue curve) decay more rapidly with pair distance than those of the unconstrained model (green curve, repeated from Fig. 5F).

### Supplementary Tables

| Crystal | Wedge | Frames | Spindle Range (deg.) | $\langle I/\sigma I \rangle$ |
|---|---|---|---|---|
| 7 | 1 | 1-500 | 5-55 | 16.0 |
| 8 | 1 | 1-500 | 300-350 | 14.0 |
| | 2 | 1-500 | 255-305 | 15.6 |
| | 3 | 1-500 | 130-180 | 14.6 |
| 9 | 1 | 1-500 | 0-50 | 18.7 |
| | 2 | 1-500 | 45-95 | 18.1 |
| 10 | 1 | 1-500 | 250-300 | 16.2 |
| | 2 | 1-500 | 295-345 | 23.1 |
| | 3 | 1-500 | 340-390 | 18.7 |
| | 6 | 1-500 | 70-120 | 20.3 |
| | 7 | 1-500 | 115-165 | 14.9 |

**Table S1:** The X-ray datasets that were combined and used for both structure determination and diffuse scattering analysis in this study.

| | PDB ID *6o2h* |
|---|---|
| **Data collection** | |
| Space group | P 1 |
| $a$, $b$, $c$ (Å) | 27.42, 32.13, 34.51 |
| $\alpha$, $\beta$, $\gamma$ (°) | 88.66, 108.46, 111.88 |
| Mosaicity (°) | 0.030 |
| Resolution range (Å) | 32.55–1.21 (1.23-1.21)Values in parentheses are for highest-resolution shell. |
| $R_{\text{pim}}$ | 0.025 (0.038) |
| $\langle I/\sigma(I)\rangle$ | 27.4 (12.5) |
| $CC_{1/2}$ | 0.995 (0.994) |
| Completeness (%) | 96.1 (52.9) |
| Multiplicity | 4.7 (1.4) |
| **Refinement** | |
| Resolution (Å) | 32.55–1.21 |
| Unique reflections: all / free | 30108 / 1495 |
| $R_{\text{work}}$/ $R_{\text{free}}$ | 0.096 / 0.117 |
| Number of non-H atoms | |
| Protein | 1122 |
| Ligand/Ion | 29 |
| Water | 92 |
| Mean isotropic B-factors | |
| Protein | 12.45 |
| Ligand/Ion | 31.44 |
| Water | 26.25 |
| Model validationCalculated using MolProbity (*44*) | |
| Ramachandran outliers (%) | 0 |
| Ramachandran favored (%) | 99.21 |
| Rotamer outliers (%) | 0.81 |
| C-$\beta$ deviations | 0 |
| R.m.s. bond lengths (Å) | 0.0272 |
| R.m.s. angles (°) | 2.31 |
| Clashscore | 0.00 |
| Overall score | 0.50 |

**Table S2:** Data collection and refinement statistics.

| Wavevector direction | Longitudinal fraction (%) | $v_s$ (m/s) |
|---|---|---|
| Γ-L | 7.2 | 601.2 |
| | 12.1 | 819.6 |
| | 99.1 | 1270.3 |
| Γ-M | 1.0 | 744.4 |
| | 18.1 | 606.1 |
| | 98.3 | 1033.4 |
| Γ-N | 20.6 | 675.7 |
| | 28.0 | 876.5 |
| | 93.6 | 1170.9 |
| Γ-R | 18.4 | 852.6 |
| | 22.8 | 769.1 |
| | 95.7 | 1202.6 |
| Γ-X | 1.7 | 694.0 |
| | 22.7 | 775.9 |
| | 97.2 | 1294.0 |
| Γ-Y | 26.9 | 708.5 |
| | 51.3 | 598.1 |
| | 81.8 | 1059.2 |
| Γ-Z | 0.9 | 693.5 |
| | 9.4 | 793.0 |
| | 99.6 | 1047.4 |

**Table S3:** Acoustic phonon properties. Wavevector directions are defined in Fig. S15. Longitudinal fraction and speed of sound ($v_s$) were calculated for each of the three acoustic phonons at a fractional wavevector magnitude of 0.1. The longitudinal fraction was defined as $(\hat{\mathbf{u}} \cdot \hat{\mathbf{k}}) \times 100\%$, where $\mathbf{k}$ is the wavevector and $\mathbf{u}$ is the principal axis of vibration of the protein center of mass.

| | |
|---|---|
| Protein center of mass (x,y,z) (Å) | (-0.943, 14.007, 24.322) |
| Center of reaction (x,y,z) (Å) | (-1.844, 13.061, 25.358) |
| $\mathbf{T}$ ($T_{1,1}$, $T_{2,2}$, $T_{3,3}$, $T_{1,2}$, $T_{1,3}$, $T_{2,3}$) (Å$^2$) | (0.0385, 0.0443, 0.0377, 0.0009, 0.0037, 0.0024) |
| $\mathbf{L}$ ($L_{1,1}$, $L_{2,2}$, $L_{3,3}$, $L_{1,2}$, $L_{1,3}$, $L_{2,3}$) (deg.$^2$) | (0.5675, 0.7941, 0.6688, -0.0789, -0.0383, -0.1438) |
| $\mathbf{S}$ ($S_{1,1}$, $S_{2,2}$, $S_{3,3}$, $S_{1,2}$, $S_{1,3}$, $S_{2,3}$) (deg.Å) | (0.0109, -0.0075, 0.0034, -0.0103, -0.0012, -0.0079) |
| r.m.s. displacement = $(\mathrm{tr}(\mathbf{T})/3)^{1/2}$ (Å) | 0.200 |
| r.m.s. rotation = $(\mathrm{tr}(\mathbf{L})/3)^{1/2}$ (deg.) | 0.823 |

**Table S4:** Translation libration screw (TLS) parameters (*42*) calculated from the refined lattice dynamics model (Equation 78). The center of reaction was chosen as the origin for the rigid group, which makes **S** symmetric and minimizes the trace of **T**.
